# A framework for quality control in quantitative proteomics

**DOI:** 10.1101/2024.04.12.589318

**Authors:** Kristine A. Tsantilas, Gennifer E. Merrihew, Julia E. Robbins, Richard S. Johnson, Jea Park, Deanna L. Plubell, Jesse D. Canterbury, Eric Huang, Michael Riffle, Vagisha Sharma, Brendan X. MacLean, Josh Eckels, Christine C. Wu, Michael S. Bereman, Sandra E. Spencer, Andrew N. Hoofnagle, Michael J. MacCoss

## Abstract

A thorough evaluation of the quality, reproducibility, and variability of bottom-up proteomics data is necessary at every stage of a workflow from planning to analysis. We share vignettes applying adaptable quality control (QC) measures to assess sample preparation, system function, and quantitative analysis. System suitability samples are repeatedly measured longitudinally with targeted methods, and we share examples where they are used on three instrument platforms to identify severe system failures and track function over months to years. Internal QCs incorporated at protein and peptide-level allow our team to assess sample preparation issues and to differentiate system failures from sample-specific issues. External QC samples prepared alongside our experimental samples are used to verify the consistency and quantitative potential of our results during batch correction and normalization before assessing biological phenotypes. We combine these controls with rapid analysis (Skyline), longitudinal QC metrics (AutoQC), and server-based data deposition (PanoramaWeb). We propose that this integrated approach to QC is a useful starting point for groups to facilitate rapid quality control assessment to ensure that valuable instrument time is used to collect the best quality data possible. Data are available on Panorama Public and on ProteomeXchange under the identifier <u>PXD051318</u>.

## Introduction

Liquid chromatography coupled to tandem mass spectrometry (LC-MS/MS) is a sensitive and powerful approach to characterize the proteome using quantitative technologies. Historically, discovery experiments prioritized detecting the most peptides and proteins in a sample. The field has moved towards more complex and scaled-up applications that prioritize throughput and quantitation. Proteomics is increasingly relevant in the clinic in pursuit of disease diagnostics and novel biomarkers, and further development of stringent quality controls is necessary to propel this quantitative work going forward^1–4^. Despite best practices and best intentions, issues in sample processing and system function will occur, but the source of an issue is not always immediately clear, and prompt identification of problems is crucial for time and cost management.

In a typical bottom-up quantitative proteomics experiment, variability can be introduced from the LC-MS system function, sample preparation, and downstream analyses. Rapid, consistent, and longitudinally tracked injections of a system suitability sample can be used to verify whether an LC-MS system is functioning adequately within desired tolerances. These runs effectively serve as a “canary in the coalmine” to provide early indications of an LC-MS system regression. Crucially, if we cannot obtain consistent peak areas, retention times, and mass accuracy, then quantitative experiments will be challenging. Beyond the system, much of the variability in quantitative studies originates from sample preparation^5^. This variability can include issues in protein extraction, digestion, and clean-up, and in sample batch effects. A strategy must be in place to assess the quality of the entire process including sample preparation, signal processing, normalization, and batch correction.

Any issues with the LC or MS impact data quality and challenge interpretation of results. If the system cannot generate reproducible measurements from equivalent quantities, any results collected thereafter are not reliable. Thus, rapid identification of these issues prior to and throughout sample data acquisition is crucial. Large experiments with runtimes on the scale of weeks to months will inevitably experience changes in instrument sensitivity or chromatography, therefore the ability to quantify the degree of change in repeated system suitability measurements throughout an experiment is useful. Early attempts to assess LC-MS systems in bottom-up proteomics focused on evaluating results such as the number of identified peptides post-database searching^6,7^. These identifications were used as a proxy for system function. However, this is complicated by the time requirements and potential variability introduced by database searches. We and others incorporated a statistical process control (SPC) into our bottom-up proteomics QC workflow (reviewed by Bramwell in 2013^8^) which often focuses on using identity-free metrics to track outputs over time relative to a baseline. The system suitability protocol developed by the Clinical Proteomic Tumor Analysis Consortium (CPTAC)^9^ to evaluate targeted proteomics assays across 11 institutions and 15 instruments used chromatographic and MS metrics including normalized peak area, chromatographic resolution, peak capacity, peak tailing, retention time drift, column conditioning and carryover. The Anubis/QCHtmlSummary platform used SRM to track longitudinal system function in peptides derived from a bovine protein mixture^10^. Our group incorporated SPC into SProCop and later automated SPC in Panorama AutoQC^11,12^. Many QC platforms have been developed in the past decade that incorporate longitudinal system monitoring and use multi-variable metrics to identify when the LC-MS system is not functioning optimally. To name just a fraction of this work spanning cloud-based applications, programming packages, or web interfaces includes SIMPATIQCO^13^, Metriculator^14^, iMonDB^15^, InSPECtor^16^, MSstatsQC^17^, QC-ART^18^, QCloud/QCloud2^19,20^, RawTools^21^, and Rapid QC-MS^22^. In addition to software tools, a stable and easy to obtain system suitability sample that can be measured repeatedly and reliably to assess system function is crucial. Examples of such samples include the NIST reference material (RM) 8323 yeast protein extract^23^, a mixture of 6 bovine proteins^10^, and commercially-available human protein extract^24^.

In addition to instrument performance, sample processing reproducibility is invaluable to track. Sample processing is inherently variable due to differences in sample collection or storage^3,25,26^, digestion conditions^27^, enzyme efficiency^28,29^, contaminants^30^, and unidentified issues with reagents or protocols. It is crucial to identify these issues as they arise and ensure that any differences observed in the results are primarily due to biological variation rather than technical variation associated with sample preparation. Controls can be incorporated into individual samples (internal QCs), or additional samples which can be prepared alongside experimental samples (external QC samples). Early areas of focus in the development of proteomic reference materials and internal QCs involved the use of isotopically-labeled peptides for quantitation of specific proteins in targeted assays such as IS-PRM and QconCAT^31–34^. Different internal QCs can be introduced into a workflow at numerous stages to analyze more specific aspects of a process. For example, exogenous proteins such as lysozyme or ovalbumin have been used as process controls to evaluate sample preparation^35,36^. More recent efforts have focused on synthetic proteins that produce consistently digested and reproducible peptides that span the chromatographic gradient. Additionally, others have developed protein internal QCs with derivative peptides that also function as indexed retention time (iRT)^37^ for scheduling or normalization such as DIGESTIF^38^ and RePLiCal^39^. Peptide internal controls added directly to samples have been used to assess system performance, normalize results, and as iRT standards^40,41^. Protein internal QCs added at the beginning of preparation allow examination of the full process from lysis to digestion, to clean-up. Peptide internal QCs added just before MS analysis can be used to continuously monitor instrument function. When protein and peptide internal QCs are used together, sample preparation issues can be distinguished from LC-MS system problems. External QC samples are pooled samples processed from the start of an experiment and carried through an entire protocol. They can identify issues in sample preparation. For example, a known mixture of phosphoproteins can be used to assess phosphopeptide enrichment efficiency^42^. Here, we describe the use of external QC samples that are prepared multiple times within a batch and across an experiment to serve as a “known unknown” and evaluate their variability as a measure of sample preparation consistency.

Finally, once the system function and sample processing are found to be within expectations, the experimental results can be confidently considered. Many analytical tools have been developed to assess sample data quality. Early examples include the NIST MSQC pipeline^43^ and QuaMeter^44^ which often relied on identity-based metrics^44^. QuaMeter and other tools later expanded into identity-independent, multivariate analyses to assess sample runs^18,45,46^. If the experiment goal is to maximize detections, identity-based metrics to determine data quality can be useful. However, in quantitative proteomics experiments, these identity-based metrics do not adequately assess the system, sample preparation, or data quality. Identifications will change depending on the database searching scheme employed. Evaluating the consistency of metrics in raw, unnormalized data across runs, batches, and experiments is of greater utility. This includes metrics such as peak area, mass accuracy, isotope distributions, and retention times of known quantities of the same analytes. These QC metrics can aid in the selection of optimum normalization and batch correction methods. Effective ways to normalize sample data after database searching have been reviewed by others^47^. Each sample matrix and MS collection scheme is unique, and the selection of normalization methods may change accordingly. For example, data-dependent acquisition methods suffer from missing data due to the irregular nature of sampling, which complicates analyses and normalization. Existing tools such as pmartR and the ProNorM workflow have been used to improve post-processing of datasets with missing data^48,49^. While we advocate for the use of peptide-level quantitation given the ambiguities associated with protein-level quantitation^50^, approaches to normalize data at the peptide and protein-level will differ. Crucially, determining whether normalization and batch correction in a data analysis pipeline has effectively reduced variance without nullifying biological differences can be challenging^51^. We use external QC samples to assess the reproducibility and consistency of sample preparation and evaluate whether the normalization and batch correction methods used have reduced variance in the data. External QC samples are standard practice in all quantitative analysis where you have a set of samples that you know the quantity of the analyte(s) being measured and the analysis of those sample(s) provide confidence that the entire quantitative workflow (i.e. sample prep, data collection, signal processing, normalization, etc.) behaved as anticipated. We suggest having two external QC samples that are prepared and analyzed multiple times in each batch (when possible). One sample can be used to aid in signal calibration and normalization^52^. The second can be used to assess the intra- and inter-batch variance, and the effect of processing steps (including normalization) used during the analysis.

Here, we present a customizable framework for QC in quantitative bottom-up proteomics experiments including practical vignettes where this workflow was used to identify and resolve sample processing and instrumentation issues. We evaluate our quantitative proteomics experiments at three levels in 3 major stages: the system, the sample, and the entire workflow (Figure 1). Three classes of QCs are implemented using three sample types: a study-independent system suitability standard, internal QCs, and external QC samples (Figure 1, Supplementary Table 1, Supplementary Discussion and Frequently Asked Questions). These controls differ in composition, purpose, and the stage they are introduced into a workflow, but all are used as “checkpoints” for quality assessment of quantitative experiments. Furthermore, we use a digital notebook to track maintenance and known issues, peaks are visualized in Skyline^53,54^, and system suitability MS data files are uploaded automatically to PanoramaWeb^55^ using Panorama AutoQC^12^. This adaptable approach to QC has enabled the quick identification of problems in an experiment, pinpoint their origin, and improve quantitative outcomes by reducing the contribution of technical variation. We present vignettes that assess diverse sample matrices including plasma, cerebrospinal fluid (CSF), yeast, brain tissue, and commercially available reagents. We also discuss criteria for the selection and optimization of QC samples and reagents.

**Figure 1:**
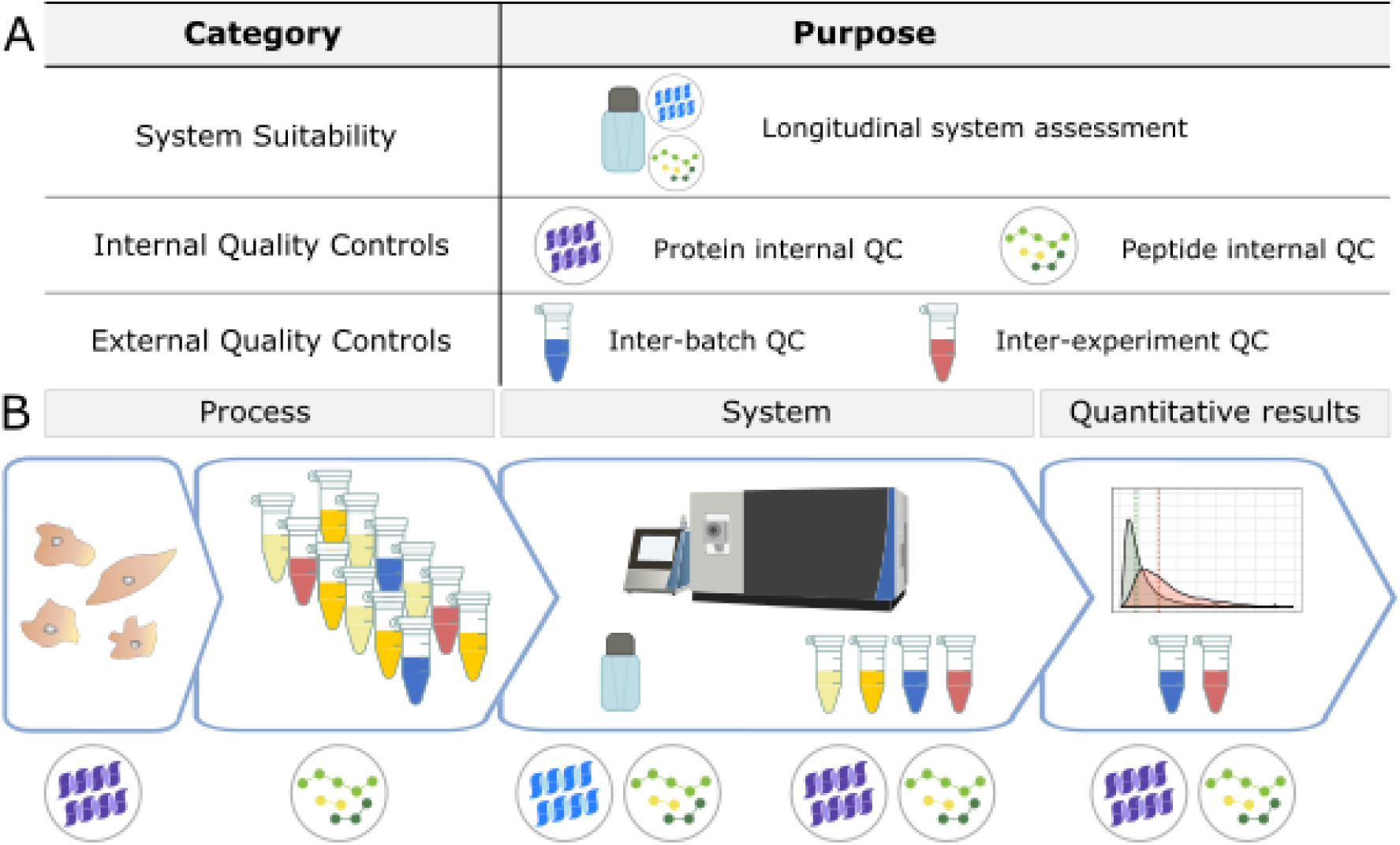
We use three categories of quality controls (QC) in our proteomics experiments: system suitability, internal QCs, and external QC samples (A). The system suitability control is used to verify that the LC-MS system is functioning before and throughout sample data acquisition. Internal QCs are added to experimental samples to assess protein and peptide-level deviations in sample processing as well as instrument function. External QC samples are additional samples prepared alongside experimental samples to monitor sample processing and batch effects. These samples are ideally formulated by pooling the experimental samples and are prepared multiple times within a batch. They also contain the same internal QCs used in experimental samples. External QC samples can serve two functions: to assess the sample preparation workflow, or to evaluate normalization methods. In the context of a workflow (B), internal and external QC samples must be planned for and incorporated beginning with sample collection and processing. Before performing any data analysis, the variance of internal and external QC samples are examined. In combination, these controls can be used to evaluate sample processing and quantitative results within an experiment, as well as the LC-MS system function during an experiment and longitudinally over years.

## Experimental Procedures

To illustrate the application of this quality control framework, we selected six vignettes that highlight the application towards system suitability, sample preparation, and evaluating the quality of quantitative data. Each example is described briefly below. All experiments with multiple batches adhered to batching and blocking procedures described previously^56,57^. More detailed Supplementary Methods are provided as part of Supplementary Material. Unless noted otherwise, system suitability and internal controls in samples and external QC samples were imported into Skyline (version 22.2) using similar settings for PRM and DIA data.

### Quality Control Sample Composition and Associated Reagents

Figure 1 describes the QCs used (A) and when they were introduced (B) into a workflow. Unless noted specifically under each section header, the following control compositions were used in the 6 vignettes we describe. For system suitability, we injected 30-150 fmol Pierce™ Retention Time Calibration (PRTC) Mixture (Thermo Scientific) in a carrier background of 600 fmol of bovine serum albumin (BSA) tryptic digest in 0.1% trifluoroacetic acid in water. The protein internal QC was 16 ng yeast enolase per µg of sample protein, which captured sample processing variation from tryptic digestion and LC-MS. The peptide internal QC (30-150 fmol per injection) of PRTC was added just prior to LC-MS analysis to capture LC-MS variation. The quantity of peptide internal QCs were kept consistent between the system suitability and experimental samples, and the peptides tracked by PRM are listed in the Supplementary Material in Table S1. We used two forms of external QC samples: one to assess sample preparation, and one for normalization or batch correction. The external QC samples were composed of two of the following options, with the best options listed first: a pool derived from representative experimental samples, additional samples from the same source as the experimental samples, or the same type of sample but from another source. The same protein and peptide internal QCs were included in the external QC samples.

### Vignette 1: System Suitability on Orbitrap Eclipse Tribrid

The system suitability standard (600 fmol BSA peptides, 150 fmol PRTC peptides per injection) was separated by high-performance liquid chromatography (HPLC) using a Thermo Easy nLC1200 coupled to a Thermo Orbitrap Eclipse Tribrid Mass Spectrometer. All LC-MS method details are included in the Supplementary Material under the heading “Supplementary Methods”. The target mass list for the system suitability methods included 17 peptides and are listed in Supplementary Table 2.

### Vignette 2: MS issue on Orbitrap Fusion

The tryptic yeast proteome was obtained from Promega, which was reduced, alkylated, and digested with trypsin according to the manufacturer’s instructions. One μg of yeast peptides were injected alongside the PRM system suitability runs described above. A Waters NanoAcquity UPLC was coupled to a Thermo Orbitrap Fusion mass spectrometer were use for DDA and PRM data generation. All LC-MS method details are included in the Supplementary Material under the heading “Supplementary Methods”.

Using a Nextflow pipeline (https://github.com/mriffle/nf-teirex-dda, revision 068f68323c9f9a181175a81ee796ef4a3373b5ed), the DDA MS data were converted to mzML with msConvert^58^, peptides identified using Comet^59,60^, version 2023.01 rev. 2 (<u>uwpr.github.io/Comet/releases/release_202301</u>), q-values and posterior error probabilities at the peptide-spectrum match (PSM) level were acquired using Percolator^61^ version 3.06 (<u>github.com/percolator/percolator/releases</u>), and uploaded to <u>Limelight</u>^62^ for visualization and dissemination. Searches were performed using a Saccharomyces cerevisiae reference proteome FASTA file (Uniprot Proteome ID: UP000002311, downloaded January 27, 2024) appended with the internal control yeast enolase 1 and a common list of contaminants generated in-house. The analyzed DDA data is accessible on Limelight (<u>Project ID = 131</u>).

### Vignette 3: Sample processing issue on Orbitrap Lumos Tribrid

Sample preparations of pooled commercial human CSF (Golden West Biologicals) included four different approaches and 8 replicates. Two were variations on the paramagnetic bead-based, single-pot solid-phase-enhanced sample preparation (SP3) that was developed and optimized by others^63–66^ (labels: 1BD and 2BD) and used MagReSyn carboxylate or hydroxyl beads (ReSyn Biosciences). The other two methods tested included S-trap column (Protifi) digestion and clean-up (label: STR), and in-solution digestion with Rapigest SF (label: ISD), and mixed-mode solid phase extraction clean-up (label: MIX). A large pool of Golden West human CSF was aliquoted into 50 μL volumes of 8 replicates for each sample preparation type.

All sample preparation and LC-MS method details are included in the Supplementary Material under the heading “Supplementary Methods”. Briefly, all CSF protein was denatured (urea, RapiGest, or SDS), reduced (dithiothreitol (DTT)), alkylated (iodoacetamide (IAA)), digested with trypsin (1:25 ratio), and subjected to sample clean-up (washes, methanol:chloroform, or mixed-mode ion exchange (MCX) columns (Waters)). Yeast enolase was spiked into all samples prior to processing at 800 ng per sample as an internal protein QC. 150 fmol PRTC was added per 1 µg of sample peptides as an internal peptide QC just before LC-MS analysis on a Thermo EASY-nLC 1200 coupled to a Thermo Orbitrap Fusion Lumos Mass Spectrometer.

### Vignette 4: LC issue on Orbitrap Eclipse Tribrid

All sample preparation and LC-MS method details are included in the Supplementary Material under the heading “Supplementary Methods”. Briefly, after measuring protein concentration with the Pierce BCA assay (Thermo Scientific), 50 μg human plasma was denatured (2% SDS), reduced (DTT), alkylated (IAA), washed, and digested with trypsin. Yeast enolase was spiked in prior to processing at 800 ng per 50 μg sample as an internal protein QC. 150 fmol PRTC was added per 1 µg of sample peptides as an internal peptide QC just before LC-MS analysis on a Thermo EASY-nLC 1200 coupled to a Thermo Eclipse Tribrid Mass Spectrometer.

### Vignette 5: Combining system suitability and internal quality controls on Orbitrap Lumos Tribrid

All sample preparation and LC-MS method details are included in the Supplementary Material under the heading “Supplementary Methods”. Briefly, mouse brain micropunches were denatured (SDS) and homogenized in a Barocycler 2320 EXT (Pressure Biosciences Inc.), protein concentration measured using the Pierce BCA assay (Thermo Scientific), and 25 μg of homogenate was reduced (DTT), alkylated (IAA), cleaned-up using S-trap columns (Protifi), and digested with trypsin (1:10 ratio). Yeast enolase was spiked in prior to processing at 400 ng per 25 μg sample as an internal protein QC. 150 fmol PRTC was added per 1 µg of sample peptides as an internal peptide QC just before LC-MS analysis on a Thermo EASY-nLC 1200 coupled to a Thermo Orbitrap Fusion Lumos Tribrid Mass Spectrometer.

### Vignette 6: Assessing quantitative results on Orbitrap Eclipse Tribrid with external quality controls

Lumbar CSF from 280 patients were divided into four major groups: 1) Healthy Control, 2) Alzheimer’s disease/mild cognitive impairment, 3) Parkinson’s disease cognitively normal and 4) Parkinson’s disease cognitively impaired. Each row of half a 96-well plate contained 10 balanced and randomized CSF samples and two external QC samples. The inter-batch QC sample was a pool of 50 patients representing all 4 groups and the other inter-experiment QC sample was a commercially available pool of CSF. These controls were processed with the samples and used to evaluate the technical precision within and between each batch prior to and following normalization and batch adjustment.

All sample preparation and LC-MS method details are included in the Supplementary Material under the heading “Supplementary Methods”. Briefly, 25 µg of CSF samples were denatured (SDS), reduced (DTT), and alkylated (IAA), before proteins were precipitated and washed on MagReSyn Hydroxyl beads. Digestion was done with trypsin (1:10 ratio). Yeast enolase was spiked in prior to processing at 400 ng per 25 μg sample as an internal protein QC. 150 fmol PRTC was added per 1 µg of sample peptides as an internal peptide QC just before LC-MS analysis on a Thermo EASY-nLC 1200 coupled to a Thermo Orbitrap Eclipse Tribrid.

We generated chromatogram libraries from each plate using 6 gas-phase fractionated, narrow-window library injections and used those runs to empirically correct a Prosit Library as described previously^67,68^. The results from this analysis were saved as a “Chromatogram Library” in EncyclopeDIA’s eLib format and used to evaluate the “wide-window” sample runs. Skyline was used to map peptides to proteins, perform peak integration, manual evaluation, and generate reports. All levels of data are available via <u>PanoramaWeb</u>.

### Data Accessibility and Figure Generation

Raw files (DIA, PRM, DDA), Skyline documents, processed results (DIA, PRM) used as input for figure generation, FASTA files, EncyclopeDIA files, metadata, and the large input files (>25 MB) for R scripts are available on PanoramaWeb (panoramaweb.org<u>/maccoss-sample-qc-system-suitability.url</u>) and registered via the ProteomeXchange with identifier <u>PXD051318</u>. The processed DDA results in Figure 3 are accessible on <u>Limelight</u>^62^ under the identifier <u>Project 131</u>. The input files <25 MB and the R code to generate Figures 3-6 (maccoss_qc_figures_3_4_5_6_S1.Rmd) and Figure 7 (plot_7a.r, plot_7b.r) are available for download on GitHub under an Apache 2.0 license: (<u>github.com/uw-maccosslab/manuscript-qc-system-suitability/</u>).

## Results

To illustrate the application of the quality control framework, we shared six vignettes encompassing the evaluation of system suitability, sample preparation, and assessing the quality of quantitative data. Each vignette represents a different experiment conducted in different sample types and on different systems, which were divided into representative, labeled method and result sub-headers.

### System Suitability

Ensuring optimal system function prior to data collection and assessment was paramount. The combined use of system suitability injections and the inclusion of internal QCs in experimental samples were useful to monitor the function of the LC-MS/MS during an experiment. Importantly - these two controls used in parallel helped instrument operators determine if changes to the system needed to be made.

The system suitability control served three functions: evaluation of the system at the start of an experiment, near real-time assessment during an experiment, and the longitudinal tracking of the system. Our system suitability LC-MS/MS runs described here used 30 minute gradients coupled with data acquisition using parallel reaction monitoring (PRM) and a sample containing 600 fmol/injection of a tryptic digest of bovine serum albumin (BSA) and 150 fmol/injection of Pierce Peptide Retention Time Calibration Mixture (PRTC) peptides. The system suitability data was automatically uploaded to PanoramaWeb^55^ using Skyline^53,54^ and the Panorama AutoQC loader^12^. This process was automated and enabled the visualization and long-term tracking of numerous QC metrics across all the laboratory LC-MS systems. Levey-Jennings plots^69^ were used to monitor individual points, while cumulative sum charts, moving range, trailing average, and trailing CV plots captured longer-term trends in the data. A subset of tracked metrics used most often include transition or precursor areas and their ratio, mass accuracy, retention time, and isotope dot product (idotp) values. In Vignette 1, the transition area and rolling CV during a typical month of system suitability runs on our Orbitrap Eclipse is shown in Figure 2A and 2B. Finally, we paired data deposition and tracking with user-provided annotations and logbooks. Anytime a new instrument operator ran an instrument, they could easily access historical data about their system to troubleshoot issues and assess how the current system performance compared to historical metrics.

**Figure 2:**
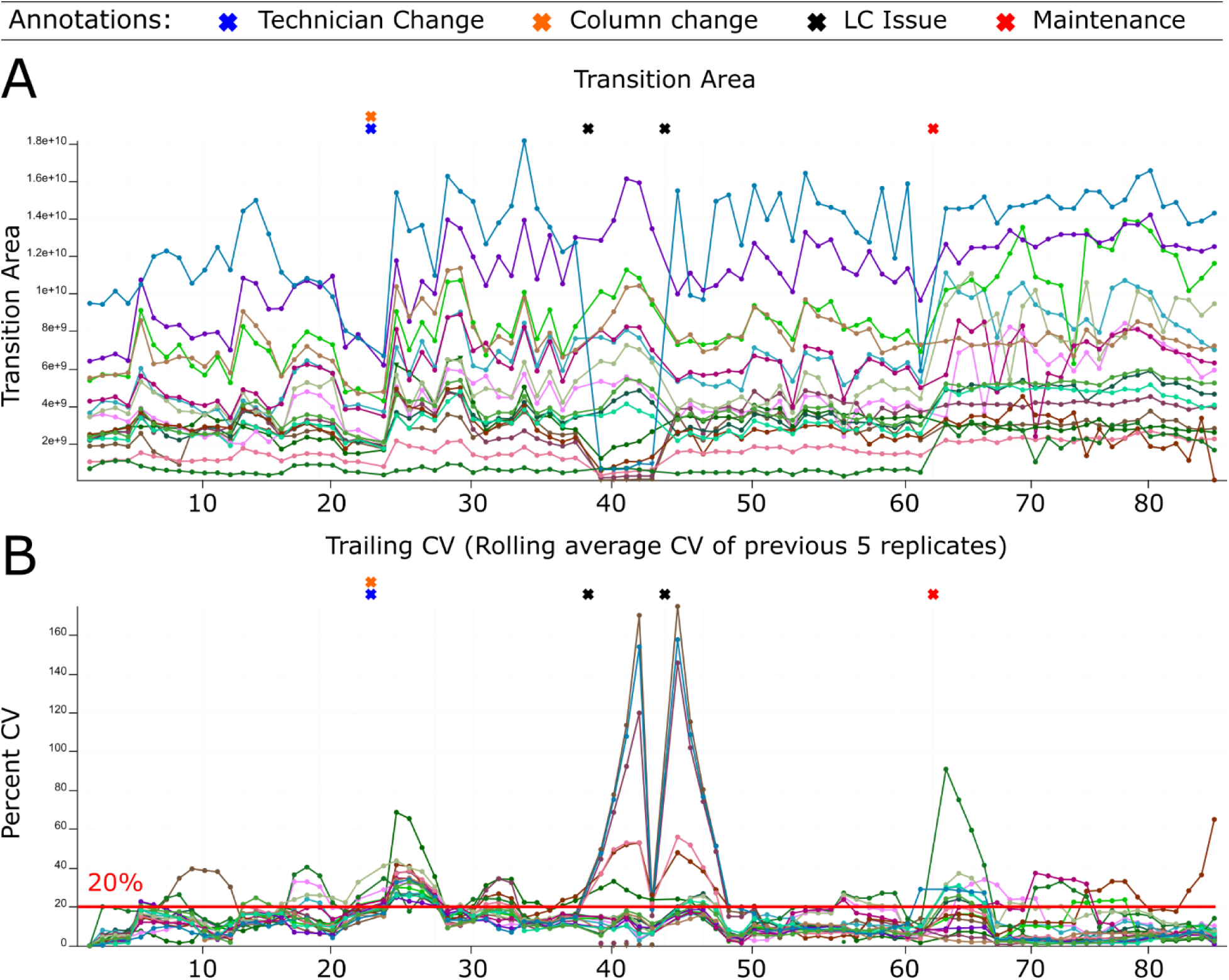
Monitoring LC-MS system performance longitudinally. System suitability tests are targeted (PRM) runs tracking the LC-MS response of consistent quantities of control peptides before, during, and after experimental runs. Raw data files are automatically uploaded and viewed online using Skyline with Panorama AutoQC. This facilitates longitudinal assessment of the system’s function via numerous metrics including precursor area, transition area, precursor/transition area ratios, retention time, mass accuracy, and more. This approach can be used to identify functional deficits that are not always apparent in the standard approach that simply monitors DDA peptide identification counts. We show system suitability runs from an Orbitrap Eclipse Tribrid from March 1, 2022 - April 10, 2022 including the instrument operator annotations and two experimental metrics: (A) the transition area of individual runs and (B) the trailing CV of the previous 5 replicates. A column and operator change occurred on March 13 (orange and blue “X”s). The transition area (A) and precursor area (not shown) dropped starting the morning of March 21st (first black “X”). The instrument operator postulated there was an issue with the LC solvents, but queued up system suitability runs to confirm the system decline. The trailing CV also clearly showed the change in the transition area (B) and precursor area (not shown). After several hours when performance had not improved, the sample pump buffer was replaced (Second black “X”). Shortly after, the system stabilized. On April 1, routine maintenance to clean the quadrupole was performed and a new set of runs were started.

In developing our approach to assessing system suitability, we shifted to using targeted (PRM) methods that track known peptides in a reasonably simple mixture. This provided a more accurate picture of the current state of a system’s function than classical approaches which consider identification-based metrics. In Vignette 2, our group had a Thermo Orbitrap Fusion coupled to a Waters NanoAcquity ultra-performance liquid chromatography (UPLC) system that exhibited inconsistency in data collection due to unknown causes from late 2014 into 2016 (Figure 3). At that time, we were using both DDA and targeted assays to assess system suitability. Historically, we and other groups would consider peptide identifications and PSMs in a commercially available or in-house complex digest measured by DDA as a fast proxy for the functioning of our MS platforms. We have since pivoted to rapid evaluation of multiple metrics of stability including the chromatography, retention time, mass accuracy, and peak intensity of a reasonably simple pooled sample generated in-house. This allowed the group to evaluate differences between instruments, discern LC versus MS issues, and identify deeper problems in the MS that may not be captured by relying on identification metrics.

**Figure 3:**
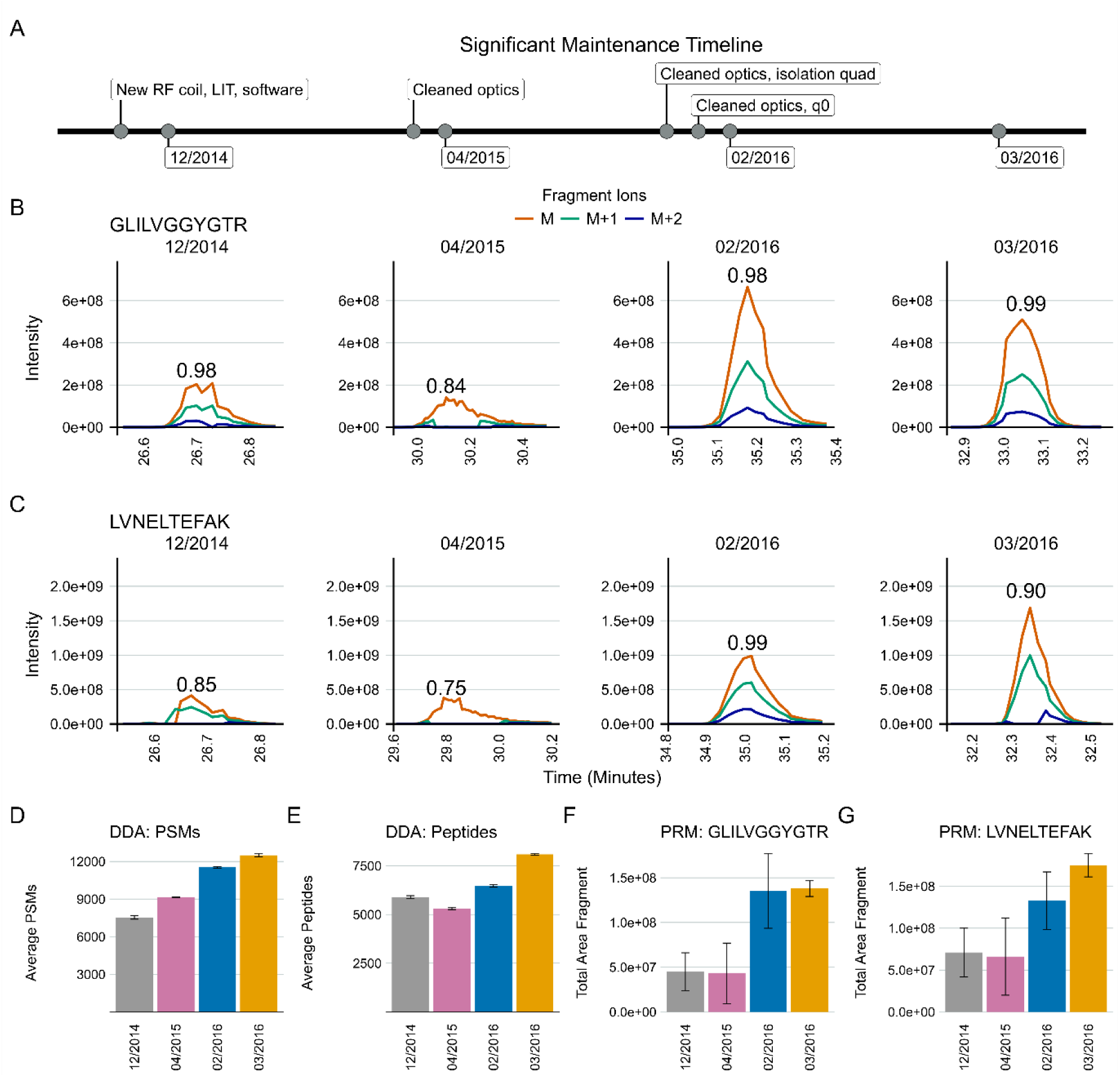
DDA spectral counting and peptide identifications fail to detect severe instrument failure. An Orbitrap Fusion exhibited inconsistent data collection for years. Typically, after cleaning various ion optics the system performance would be evaluated using DDA spectral counting and targeted PRM methods. Three different sessions spanning approximately 16 months from this time are shown to illustrate why targeted (PRM) system suitability runs were adopted. Major periods of maintenance in close proximity to these runs are shown in (A). We frequently, but inconsistently, observed the loss of ^13^C isotope peaks M+1 and M+2 precursors in the MS1 chromatograms of peptides in our PRM system suitability tests. The idotp values are printed on the chromatograms. Many peptides exhibited these issues during this time, but one representative replicate of GLILVGGYGTR (B) and LVNELTEFAK (C) are shown for brevity. Cleaning the optics in April 2015 only seemed to make the PRM chromatograms (B and C, second panels) worse. At the same time, DDA-identified PSMs increased (D) and identified peptides decreased slightly (E). The total area fragment of GLILVGGYGTR (F) and LVNELTEFAK (G) did not improve. In 2016, additional ion optics were cleaned (A). Most notable was some build up on the bent quad (q0) that was removed. At that point, the system began to improve, and we saw signal intensities improve in the 2016 batches in the DDA (D) and PRM (B-G) runs. However, it was clear that the underlying hardware issue had not been resolved as we still observed inconsistent, seemingly random loss of M+1 and M+2 signal in 2016 while the DDA-derived PSMs and peptide identifications continued to climb, suggesting these approaches do not serve to capture even significant instrumentation issues.

A standardized yeast proteome digest was measured with DDA across three months - December 2014, April 2015, February 2016, and March 2016. Within the same instrument runs, we measured the mixture we currently use for system suitability with PRM (600 fmol BSA, 150 fmol PRTC per injection). A timeline of significant maintenance on the MS during this time is provided (Figure 3A). Representative examples of PRM precursor chromatograms are shown for GISNEGQNASIK (PRTC, Figure 3B) and for LVNELTEFAK (BSA, Figure 3C) in each dataset. We frequently, but not predictably, observed a loss of M+1 and M+2 ^13^C isotope peaks in precursor chromatograms in system suitability runs beginning in 2014 (Figure 3B and 3C, left panel). This was reflected in idotp values (shown above chromatographic peaks). By 2015, after cleaning the front optics, the performance declined even further (Figure 3B and 3C, second panels). The total area fragment intensities of GLILVGGYGTR (F) and LVNELTEFAK (G) did not increase substantially. However, during this transition, the number of PSMs increased despite the worse instrument function. Peptide identifications by DDA did nominally decline, although the decrease was not commensurate with the degree of instrumentation failure. Later in 2016, additional ion optics were cleaned (A) including visible build-up on the bent quad (q0) which was removed. Signal intensities improved markedly in terms of both the DDA (D) and PRM (B-G) runs. However, it was evident that additional hardware issues were present since we still continued to observe sporadic loss of M+1 and M+2 signal in the two 2016 batches shown here. Concurrently, the PSMs and peptide identifications in the DDA runs continued to increase despite clear instrumentation issues.

### Internal quality controls to assess sample preparation

Internal QCs were added to all experimental samples to assess protein digestion, peptide recovery, and to distinguish between sample preparation issues and post-digestion measurement problems. We most often used yeast enolase 1 (ENO) protein as a protein internal QC to monitor digestion and peptide recovery when working with mammalian samples. PRTC was spiked into samples after digestion and clean-up just before injection as a peptide internal QC.

In Vignette 3, we used protein and peptide internal QCs to troubleshoot while standardizing a digestion and clean-up protocol in human cerebrospinal fluid (CSF). We incorporated the same protein (ENO) and peptide (PRTC) internal QCs in different preparations of pooled CSF to enable more direct comparison between the methods tested. Four approaches were examined: two different functionalized magnetic bead preparations based on in series-digestion with protein aggregation (labels: 1BD and 2BD), S-trap digestion and clean-up (label: STR), and in-solution digestion with Rapigest SF (label: ISD) with MCX clean-up. The digestions were performed using the same pool of human CSF and run on a Thermo Orbitrap Fusion Lumos Mass Spectrometer coupled to a Thermo Easy-nLC with nanoflow chromatography.

Throughout the course of running these samples, the ISD condition had significantly reduced ENO peptide abundance. However, the PRTC peptides were consistent with the other protocols. The system suitability injections interspersed between sample runs showed no significant changes (not shown), which suggested the problem was protocol-specific rather than a problem with the LC-MS/MS system. The instrument operator initially postulated that they had made a mistake in pipetting a crucial reagent during the ISD preparation. To confirm the performance of the four methods without ambiguity, the entire experiment was repeated with additional caution and 8 replicates of pooled CSF. The same results were observed as in the first replicate experiment described above, and a representative example is shown for the ENO peptide VNQIGTLSESIK in Figure 4. Supplemental Figure 1 includes an additional ENO peptide (AADALLLK, A and C) and two PRTC peptides (NGFILDGFPR, E and G; TASEFDSAIAQDK, F and H). Despite the abundance of caution taken by the instrument operator, a number of runs were found to be missing appreciable levels of ENO peptides (Figure 4A and Supplemental Figure 1, A and B), but contained expected levels of PRTC (Supplemental Figure 1, E and F). When the data were re-ordered by sample group, it became clear that the ENO peptides in the ISD prep were consistently absent (Figure 4B and Supplemental Figure 1, C and D) and PRTC peptides were congruent with the other 3 protocols (Supplemental Figure 1, G and H). No significant deviations were seen in the system suitability injections throughout the run (not shown). Many of the reagents between the different preparations were shared including dithiothreitol, iodoacetamide, trypsin, ENO, and PRTC. We postulated the only reagent used exclusively in the ISD preparation was likely the source of the problem. Upon further inspection, the stock solution of 0.2% Rapigest SF (Waters) in water had expired. We postulated that an incomplete denaturation could hinder digestion and explain the lack of ENO peptides while PRTC peptides, which were added after digestion and clean-up, would perform as expected.

**Figure 4:**
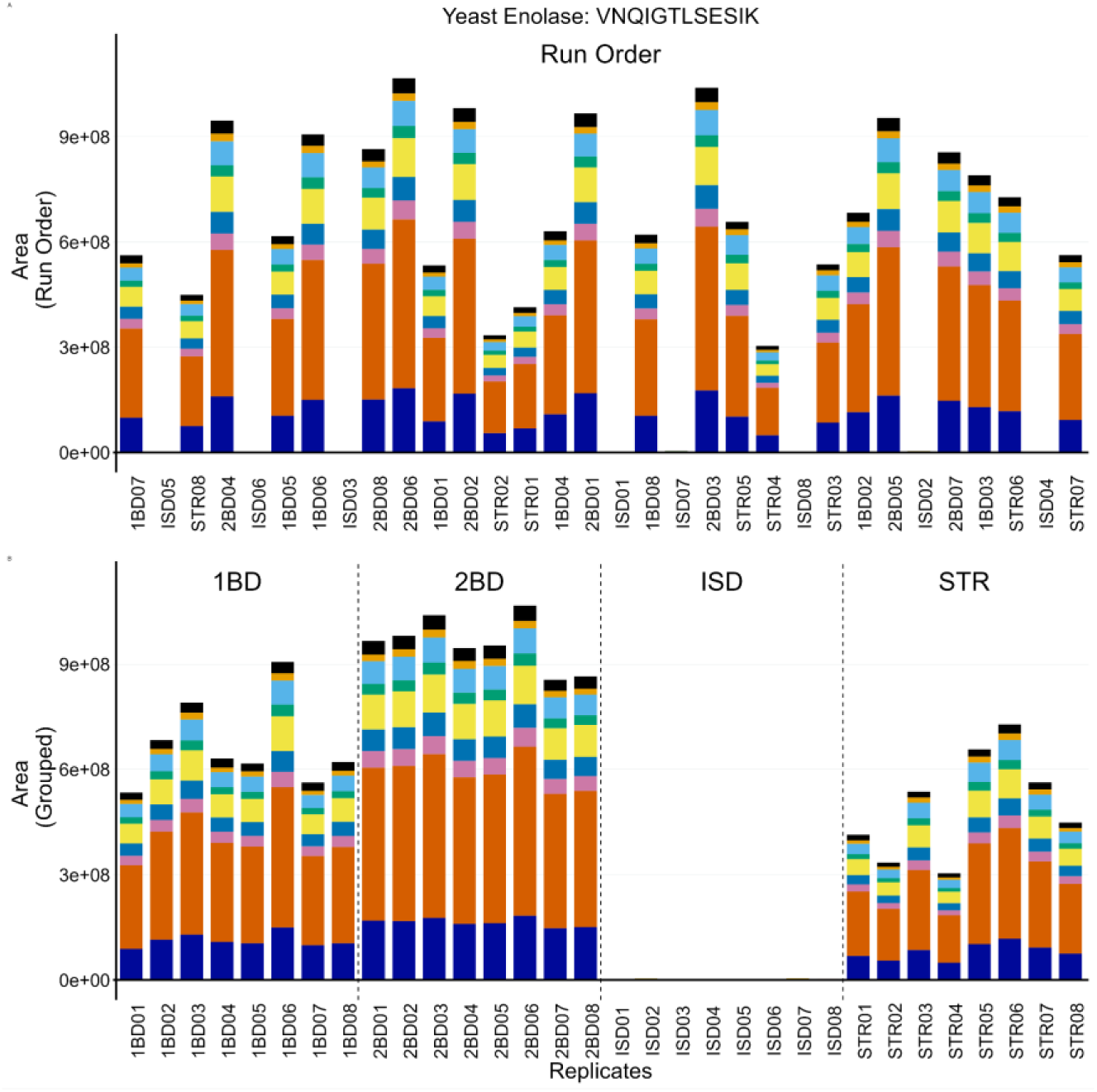
Internal QCs added to experimental samples are used to assess the entire process from lysis to LC-MS. Four different sample preparation protocols were tested - two SP3 variants (1BD and 2BD), S-trap (STR), and in-solution (ISD). All the samples were spiked with ENO as the protein internal QC to evaluate the overall processing. The transition peak area of VNQIGTLSESIK is shown here. The run order for the 8 replicates from each condition were randomized, as shown in panel (A). At inconsistent intervals, numerous samples were missing ENO peptides, whereas the PRTC peptides looked normal (Supplementary Figure 1). The cause of the missing enolase peptides became clear when the samples were grouped by sample preparation (B), revealing that there was an issue with the ISD sample preparation method. We later found that our Rapigest stock, which was one of the only reagents different between the four protocols, was expired and could have hindered denaturation, and thus digestion, in the ISD samples.

### Higher temporal resolution measurement of system state using internal quality controls

We have found that internal QCs in our sample injections can also serve as a measure of system functionality. While our system suitability injections are frequently performed every 8 to 12 hours, depending on the instrument operator and the experiment in question, the consistency of protein and peptide internal QC peptides can be examined after every sample injection to monitor issues as they arise.

Spontaneous issues with sample injections and the samples themselves happen. We share one such case in Vignette 4 from a Thermo Easy-nLC with nanoflow chromatography coupled to a Thermo Orbitrap Eclipse Mass Spectrometer. While running a series of human plasma samples, we identified a bad injection using the protein (ENO) and peptide (PRTC) internal QCs present in the samples. The problematic injection is denoted “7” in Figure 5. The peak areas of ENO (Figure 5A) and PRTC (Figure 5B) were reduced relative to the other samples, and the peak shape was markedly worse (Figure 5C and E).

**Figure 5:**
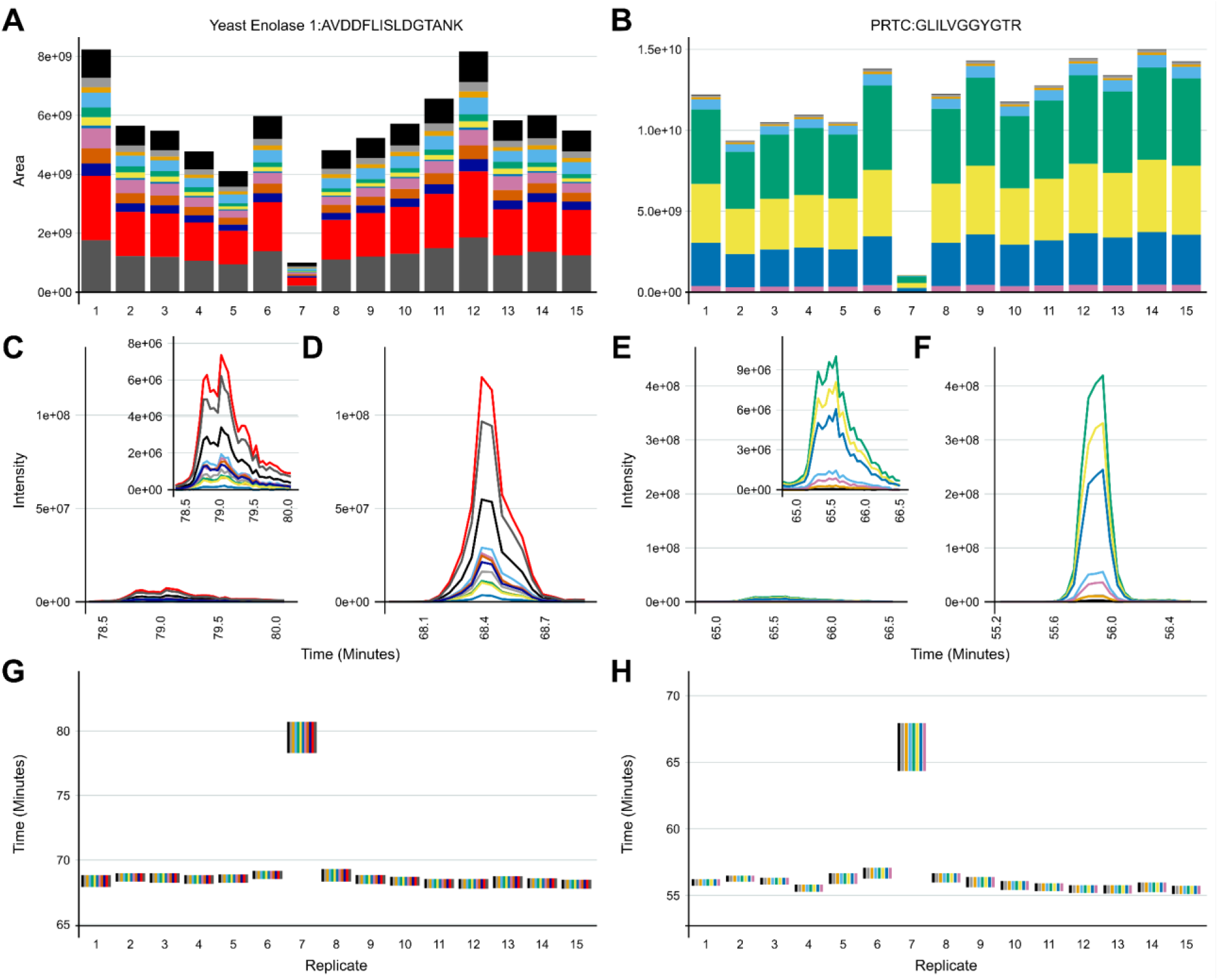
The two internal QCs can help distinguish between sample preparation versus LC-MS problems. In a series of plasma samples analyzed on an Eclipse Tribrid, one injection denoted 7 was found to have a reduced peak intensity and altered retention time across all of the internal QC peptides. One peptide from each control is shown here for brevity. In ENO, AVDDFLISLDGTANK transition areas were reduced (A), chromatography was poor (C), and the retention time was shifted (G). GLILVGGYGTR from PRTC shared a reduction in peak area (B), poor chromatography (E), and shifted retention time (H). Upon manual inspection of the LC after injection 7, the instrument operator determined that the outlet line coming from the injection valve had partially clogged. Once the repair was completed, three system suitability injections (not shown) were found to be stable and with comparable signal intensity to before the clogged line. Another sample was injected (08) and then the previous sample was reinjected into the system and is shown as 9. The transition peak areas and chromatography of AVDDFLISLDGTANK (A, D) and GLILVGGYGTR (B, F) were normal relative to the previous samples, and the retention times were again in line with other samples (G, H).

Additionally, their retention times were shifted later by several minutes (Figure 5G and H). Right after sample 7 was run, manual inspection of the LC lines by the instrument operator revealed a clogged outlet line coming from the sample injection valve. Once this was replaced, sample 7 was reinjected into the system and is denoted as sample 9. The peak areas (Figure 5A and B), peak shape (Figure 5D and F), and retention times (Figure 5G and H) in ENO and PRTC peptides were now consistent with the previous sample runs.

### Combining internal quality controls and system suitability

One might question whether it is worth the time collecting the system suitability data when the same PRTC peptides are spiked into each sample as an internal QC. However, we would argue that it is important to periodically evaluate instrument performance without the complications of varying sample matrices, sample preparation methods, or instrument operators. The combined use of system suitability injections and samples containing protein and peptide internal QCs allowed the instrument operator to identify a significant system issue in Vignette 5.

Suspected charging was observed on an Thermo Orbitrap Fusion Lumos Mass Spectrometer coupled to a Thermo Easy-nLC 1200 with nanoflow chromatography. We spent approximately 5 months troubleshooting inconsistent system performance after a turbo pump exploded while the MS was running under vacuum, which dispersed debris into the system. Key maintenance that occurred during this time are shown in Figure 6A and Supplementary Figure S2A in relation to a series of system suitability and sample runs from four instrument runs spanning about 4 months. The sample injections were all mouse brain peptides acquired using DIA in 110-minute gradient runs. Eight PRTC peptides are shown in Figure 6B. The runs are broken down by system suitability runs (SS, white background), and internal QCs from sample runs (IQC, grey background). To simplify the figure, only a subset of all the runs from August-December are shown. The runs are representative of observed trends in the full dataset, which is available on PanoramaWeb for PRTC, ENO, and BSA peptides.

**Figure 6:**
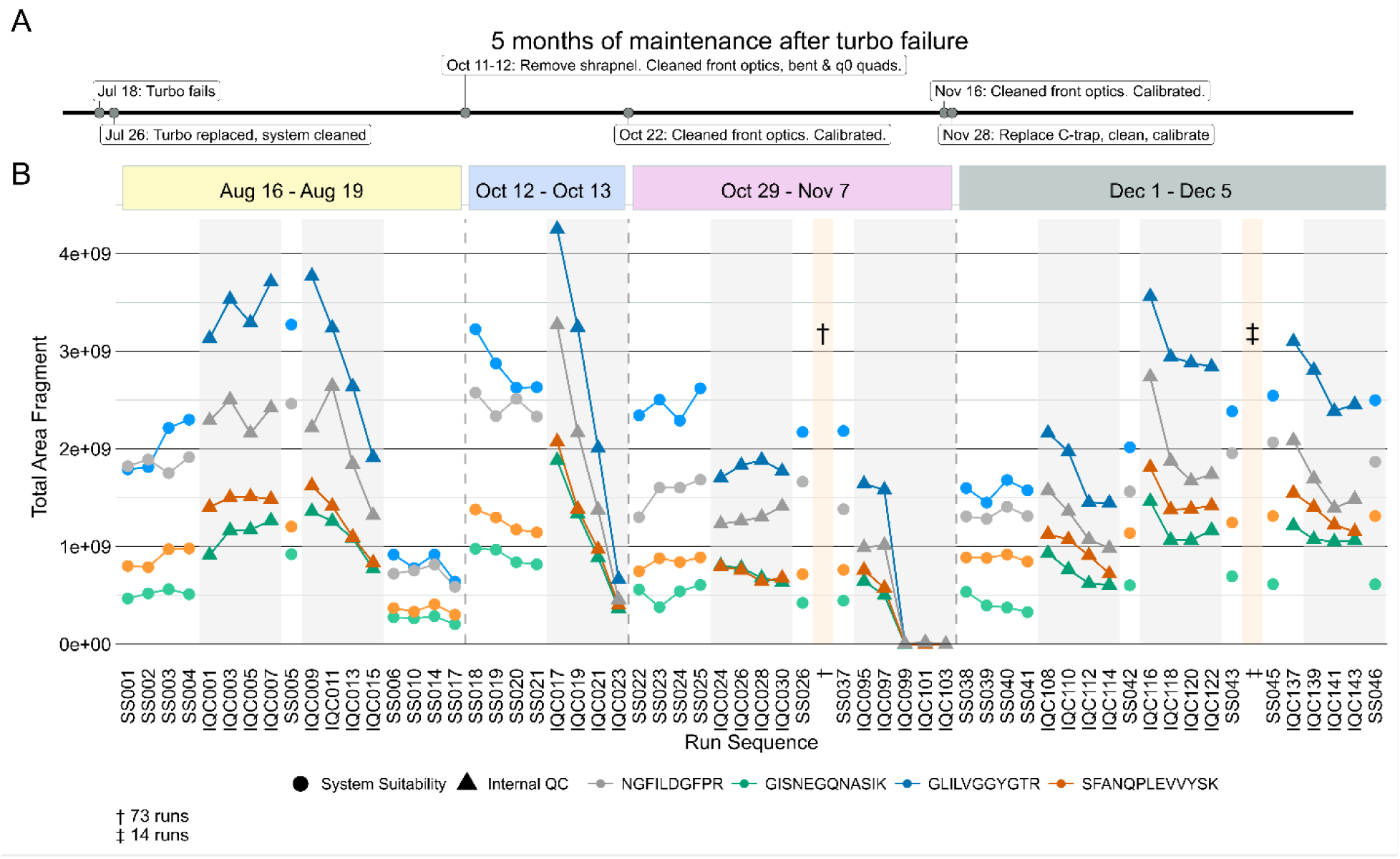
Combining system suitability injections and sample internal QCs facilitate real-time monitoring of the system. On an Orbitrap Lumos Tribrid, a significant turbo failure led to months of inconsistent and unpredictable data collection. A rough summary of the major maintenance steps during 5 months of troubleshooting post-turbo failure are shown here (A). Between August - December, signal intensity would unpredictably begin decreasing with each injection of two different run types. Four PRTC peptides from system suitability runs (B, white background) and sample internal QCs (B, grey background) illustrate this in four different experimental batches. Only a subset of all runs are shown here to improve visibility, but the trends are representative. The full dataset including all runs for PRTC, ENO, and BSA are available on Panorama. In August, after 7 sample injections (IQC001-IQC007) and 5 system suitability injections (SS001-SS005) the system was found to be stable. As additional samples were run (IQC009-IQC015), the signal intensity dropped significantly. This reduced performance was confirmed with system suitability injections (SS006-SS017). The system was taken offline. Additional metal debris from the turbo failure that was missed in earlier maintenance was removed, and the front optics and quads were cleaned. After calibration, the same issue of rapidly declining signal was observed in mid-October (SS018-IQC023). After taking the system down and cleaning the optics again, the system seemed stable for 75 runs (SS026-SS037) until the signal intensity declined rapidly while running samples IQC097-IQC103. The system was taken down again. In late November metal debris was found lodged in the C-trap, and it was replaced along with another thorough cleaning. We then observed a return to expected signal stability and intensity (SS038 through SS046) relative to matched sample batches run prior to the system failing.

After the turbo pump failed, the MS was stopped, and the internal components cleaned thoroughly. After passing vendor calibration and other in-house assessments, the experimental runs were restarted. Although the system was initially consistent (SS001-SS005 and IQC001-IQC007), the peak areas of PRTC and ENO (not shown) in sample injections started to decrease rapidly (IQC009-IQC015) with each injection. The subsequent system suitability injections (SS06-SS17) queued by the instrument operator confirmed a loss of system function relative to baseline. The system was taken offline (first dashed line). More metal debris was found and removed, and the front optics, bent quad, and q_0_ were cleaned. After this maintenance and recalibration, the system suitability runs were consistent at the start of this second run (SS018-SS021), but PRTC and ENO (not shown) declined quickly during the first set of sample runs (IQC017-IQC023). The system was taken offline again, and additional maintenance was performed including further searches for debris and cleaning the optics followed by re-calibration (second dashed line). The same issue occurred near the end of a longer run (SS022-IQC103) where 73 intermediate, stable runs are denoted by “†”. The final sample injections exhibited a loss in sensitivity. The run was halted, and the system was taken offline. Upon closer inspection of the instrument interior, additional metal debris was found lodged inside of the C-trap, and it was replaced (third dashed line). The system was stably restored to normal function. The same samples which had declining sensitivity from August 16 through November 7 were reinjected and did not exhibit the same behaviors here. The system suitability and samples ran in December (SS038-SS046) reflected historical performance for that instrument in terms of the system suitability runs and the other sample batches from the study.

### Assessing Quantitative Results

Once the process and system were verified to have functioned as expected, we evaluated our quantitative results using a combination of internal and external QC samples. The inter-batch external QC sample was an additional pooled sample of the same matrix that is processed to evaluate sample preparation and variability within a batch. Typically, this QC sample was generated by collecting additional representative sample dedicated to the purpose of serving as a control or by pooling small quantities of representative experimental samples. The inter-experiment external QC sample, composed of the same or similar matrix as the sample, was used to assess CV across experiments. Both external QC sample pools were prepared and aliquoted before beginning the experiment and then processed in parallel with the samples. This experiment also included batching and blocking of samples based on published methods^56,57^ in consideration of the sample metadata. The internal protein control in these external QC samples can be used to evaluate how consistent sample preparation was within a batch and across batches (not shown). We share a final practical example, Vignette 6, where these controls were prepared, integrated, and applied in a human CSF study. The inter-batch external QC sample was created by pooling 50 human CSF patient samples from Healthy Controls, Alzheimer’s disease/mild cognitive impairment, Parkinson’s disease cognitively normal, and Parkinson’s disease cognitively impaired. The inter-experiment external QC sample was generated using a commercially acquired CSF stock. Both external QC samples were included to assess the inter-plate and intra-plate CVs after median normalization and batch adjustment.

We used the inter-batch external QC samples to evaluate the variation and impact of normalization within and between 8 sample batches (Figure 7A). These data were examined in series beginning with level 1 (un-normalized raw peptides), level 2 (log_2_ transformed and median-normalized peptides), and level 3 (normalized and batch-corrected data) at the peptide and protein level. The coefficient of variation (CV) calculated for all peptides in the inter-batch QC improved 6.3% after normalization and batch adjustment at the peptide-level and 16.9% at the protein level. We showed that the batches were similar through our entire analytical pipeline and that overall variability was reduced with normalization. The inter-experiment external QC samples were used to examine the effect of normalization and batch correction on peptide and protein coefficient of variance in two ways. First, by examining the relationship between coefficient of variation and the log2 median abundance (Figure 7B, left panel) with a LOESS fit of a contoured density plot (red line), and second by examining the median coefficient of variation (η, denoted with red dashed line) of relative to median abundance (Figure 7B, right panel). The raw unnormalized data, the log2 transformed and median-normalized data, and batch corrected data are available on Panorama and GitHub.

**Figure 7:**
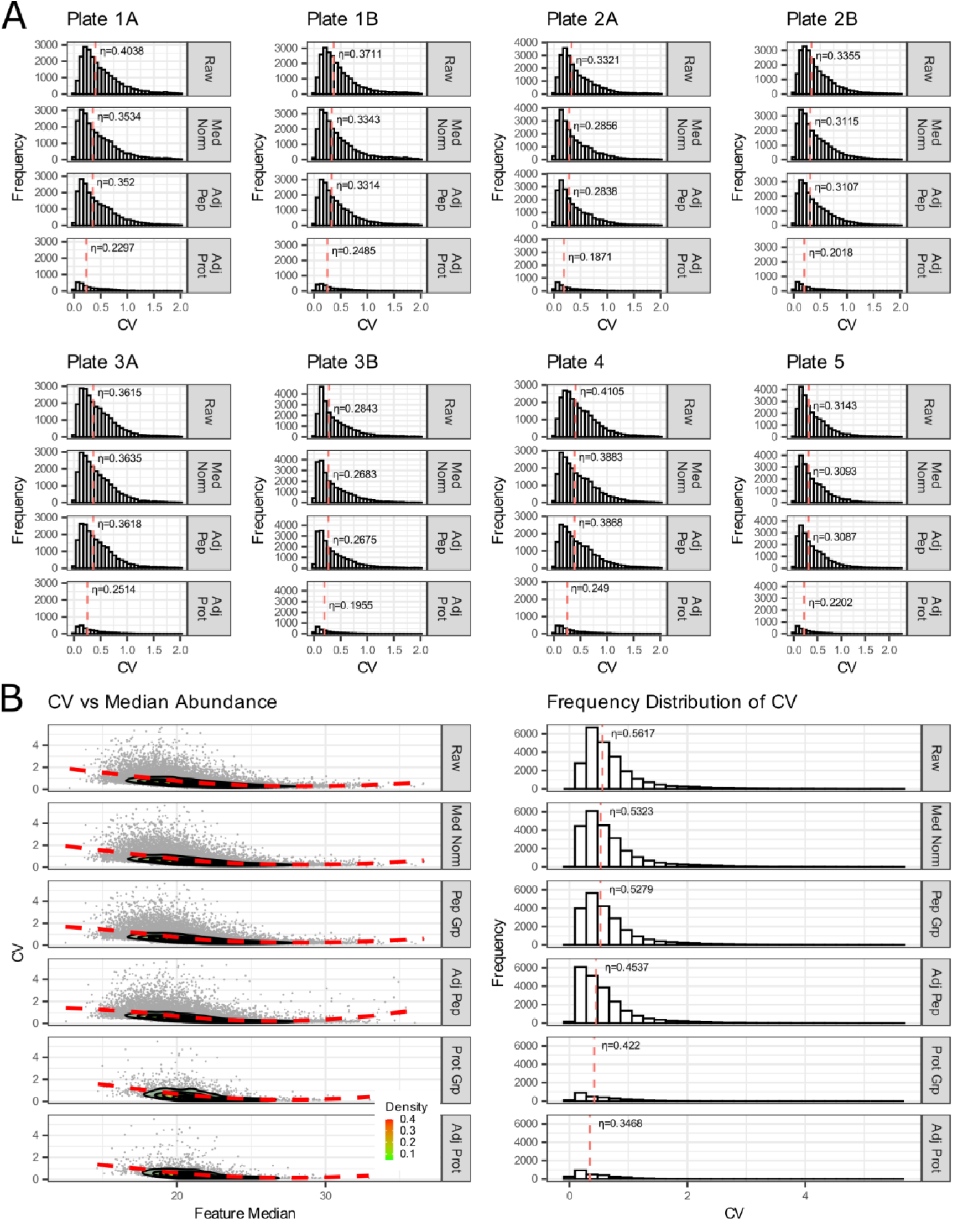
Assessment of quantitative results using median normalization and batch correction of inter-batch external QC samples. (Top panel or 7A) The effect of median normalization and batch correction of peptide and protein coefficient of variance (CV) from all the pooled lumbar CSF inter-batch external QC samples within each plate (batch). The median coefficient of variation (η) is indicated by the dashed red line. (Bottom panel or 7B) The effect of median normalization and batch correction of peptide and protein coefficient of variance from all the pooled lumbar CSF inter-batch external QC samples from all plates. The relationship between coefficient of variation and the log2 median abundance is visualized with a LOESS fit of a contoured density plot (red line).

## Discussion

Here we described six vignettes of a QC framework that incorporates long-term monitoring of LC-MS systems, the use of QCs to identify issues in sample processing and system function, and how designing experiments with integrated internal and external QC samples can improve confidence in quantitative analyses. While our group has used this QC framework in various stages for years, the series of case studies presented here represents the first detailed summary of our current QC process. Although we initially used DDA and spectral counting for system suitability measurements, we quickly moved on to targeted methods^70^, added internal QCs^71^, and incorporated external QC samples alongside internal QCs and system suitability^72^. Examples of studies where this QC workflow was applied are listed on Panorama under the “<u>Reference Information</u>” folder. This QC framework can quickly and effectively identify issues in studies of basic biology, clinical research, and in production quantitative proteomics work. All data and analyses described in this work are available on <u>PanoramaWeb</u> or <u>GitHub</u>. A more detailed summary of the QCs described in this framework, the workflow steps they evaluate, and how they were developed and optimized are listed in the Supplementary Material in Table S2 and in the “Supplementary Discussion and Frequently Asked Questions” section.

Our system suitability approach combines a reasonably simple, stable, and well-characterized reference sample, targeted MS runs before, during, and after experimental samples, and near real-time evaluation. These runs have been standardized between LC-MS instrument platforms and are longitudinally tracked with cloud-based upload and backup facilitated by PanoramaWeb and AutoQC. Using Skyline, Panorama, and AutoQC Loader, an instrument operator can track numerous metrics and provide annotations related to precursor and/or transition peak areas and ratios, retention times, isotope dot products, mass accuracy, and additional metrics. If the system suitability runs indicate both the LC and MS are functioning consistently based on the aforementioned metrics - and historical data, when available - the instrument operator can begin running experimental samples. The initial system suitability runs serve as a baseline to monitor the system throughout a run and help identify issues more quickly as they arise. We presented vignettes where system suitability runs were used to aid instrument operators in identifying problems with the LC system (Figure 2 and 5) and with the MS (Figure 3 and 6).

We use targeted runs to assess system suitability because of the rapid and straightforward analysis of targeted data in Skyline integrated with Panorama. The PRM and DDA data shown in Figure 3 indicate that DDA data does not adequately describe system suitability, which provides additional confidence in relying on a targeted approach to assessing system suitability. An Orbitrap Fusion exhibited frustratingly inconsistent loss of M+1 and M+2 precursor ions that took years to resolve because PSM number or peptide identifications in DDA runs or solely precursor and fragment ion intensities was not adequate to describe the severity of the instrumentation failure that was occurring. We show here that PSMs identified in DDA runs are decoupled from system performance as they increased even when the system was performing at its worst in 2015 (Figure 3D). Metrics including PSMs and peptide identifications derived from DDA are also subject to changes throughout the data analysis pipeline. These changes in identification metrics can include updates in the database search software or the search parameters, making longitudinal system suitability assessment particularly challenging. We propose that the use of PSMs or peptide identifications cannot always capture even significant instrumentation failures, and thus should not be relied upon to assess system suitability alone. Only through a more thorough targeted analysis of the system suitability by evaluating chromatograms, precursor and fragment intensities, mass error, and retention time stability is it possible to clearly identify system suitability issues. Towards that end, to assess system suitability, we suggest a targeted method that is applied longitudinally, repeatedly, and consistently across instrument platforms. We suggest the use of a reasonably complex peptide mixture derived from digested commercial proteins and synthetic peptides that span the gradient. Historically, we use BSA and PRTC, but reasonably many alternative proteins or synthetic peptides would be viable alternatives. Importantly, using the same peptides at the same concentration per injection that are spiked into experimental samples as an internal peptide QC is highly recommended to serve as a relative baseline of system function between system suitability runs.

The importance of protein and peptide internal QCs was highlighted in Figures 4, 5, and 6. When selecting internal QCs for a workflow, we strongly recommend assessing peptide stability in an autosampler, peptide chromatographic consistency, and signal intensity. For protein internal QCs, we also suggest evaluating digestion reproducibility. Internal QC peptides are intended for broad, lab-wide use, and testing the reproducibility and stability of potential peptides in multiple sample matrices is strongly recommended. Optimization experiments to find an ideal concentration that reliably reflects instrument performance and balances reagent cost is advised. Additionally, we suggest evaluating the performance of internal QCs when moving between reagent lots to minimize ambiguity in performance assessment. The vignettes we highlighted here show how internal protein and peptide QCs can be used to differentiate between failed sample preparation (Figure 4), LC issues (Figure 5) or MS issues (Figure 6). Including both internal QCs in all experimental samples can be used to differentiate between a single bad injection, instrumentation failures, and a systemic problem in sample preparation and give more confidence in the results of an experiment.

Protein internal QCs are added to experimental samples early in the sample preparation protocol to examine the consistency and efficacy of sample and data processing. Heavy-labeled proteins or proteins from distant species are suitable after testing their digestion efficiency, chromatographic stability, and sequence specificity in the matrix of interest. In earlier studies, we used ^15^N-labeled Apolipoprotein A1 as a protein internal QC as it was commercially available, digested reproducibly, and produces reproducible peptides that had amphipathic properties^73^. We found through internal testing that yeast enolase 1 (ENO) digested and chromatographed consistently, the peptide sequences were distinct from human enolase, and it was less expensive to use in every sample in our lab. We now routinely use yeast ENO as a protein internal QC and PRTC as a peptide internal QC. However, if the experiment involves yeast or a closely related organism, the enolase peptide signal would suffer from interference of endogenous peptides with conserved sequence and thus nullify its use as a QC. Although not discussed here, exogenous protein and peptides can also be used to monitor post-translational modification (PTM) enrichment. We have used bovine beta-casein protein coupled with ENO as a first step to distinguish between digestion deficiencies and issues with phosphopeptide enrichment. Additionally, the inclusion of heavy and light-labeled phosphopeptides spiked into a sample and described elsewhere^74,75^ can provide an estimate of the phosphopeptide enrichment efficiency.

Peptide internal QCs should be added to all samples in equivalent quantities in addition to the protein internal QCs. We prefer to use stable, standard peptides that are known to perform well on our systems. Historically, we have used a commercially available heavy peptide mixture (PRTC). However this could also be synthetic peptides without sequence overlap to the analytes of interest^37,76^ or other commercially available peptide standards such as Biognosis iRT. The quantity of PRTC added has fluctuated with time, but most frequently was 150 fmol of PRTC peptides per sample and system suitability injection on nanoflow LC systems. We have found that as low as 15 fmol PRTC/injection is sufficient on most platforms and more cost effective.

In Figure 6, both the system suitability runs and internal QCs in samples indicated the presence of a serious but intermittent LC-MS system issue. Following a catastrophic turbo pump failure, significant clean-up and troubleshooting of the MS was necessary. Simple calibration of the MS was insufficient - venting the system, disassembling, and cleaning the optics were necessary on several occasions to remove metal debris. Using the longitudinal data from our system suitability runs based on well-characterized pooled sample, we can capture intermittent problems and determine with better confidence what problems necessitate this kind of drastic maintenance rather than a standard re-calibration and heated transfer tube replacement. Finally, the inclusion of the same internal peptide QCs in the experimental samples at equivalent quantities to the system suitability sample is advised to increase the frequency that the system performance is measured. This will enable instrument operators to more quickly make changes to the system as needed.

Finally, the use of external QC samples was highlighted in Figure 7. External QC samples are two additional samples that are prepared and processed alongside experimental samples within each batch. The inter-batch QC may be used to determine whether sample preparation is reproducible between sample batches and to assess CVs. The inter-experiment QC may be used to assess the validity of the normalization and batch correction pipeline. We previously described the use of these controls with median, Log_2_ normalization, and batch adjustment of peptide signal in a recent quantitative proteomics dataset focused on the phenotypic assessment of Alzheimer’s disease in CSF^72^ in additional studies of aging and disease listed on PanoramaWeb in the “<u>Reference Information</u>” section. In recent years, we have most often used median normalization at the peptide-level^77^ but have also examined maxLFQ^78^ and directLFQ^79^ for normalization of peptide and protein-level data. Although originally developed for RNA datasets, we have found, as other groups have indicated, that linear models for microarrays (LIMMA)^80,81^ and ComBat^82,83^ reduce the contribution of technical variation in proteomics datasets. Determination of the optimum normalization and batch correction approach will likely vary according to the sample matrix and experimental approach. The use of external QC samples alongside experimental samples can provide increased confidence in selecting a final analytical approach and rigorously assessing the quantitative accuracy of the data.

Naturally, the use of significantly different samples as matrices for external QC samples will be of limited utility in assessing sample preparation. To derive the most useful inter-batch QC samples, whenever feasible, equivalent amounts of experimental samples are pooled, aliquoted, and frozen in advance to serve as the inter-batch QC. If pooling small quantities of experimental samples is not feasible, it is advised to collect additional material of the same sample matrix and generate frozen aliquots to be used solely as a control. These pooled aliquots are processed alongside experimental samples in each row of a plate, or once within a batch, and their CVs are calculated to evaluate the reproducibility of the sample preparation within that batch. The inter-experiment QC should be of the same or a very similar matrix to the experimental samples and includes protein and peptide internal QCs. This control should be aliquoted and prepared within each batch either in every row of a plate or once within a batch to assess CV.

While external QC samples are most useful in large-scale sample preparation to identify batch effects and normalize the data, external QC samples can also be useful in situations where experimental plans change unexpectedly. For example, smaller-scale pilot experiments that include external QC samples can be used to later assess how comparable a larger second, third, or fourth batch of similar samples are to one another relative to the initial experiment. They may also be useful to improve power calculations to determine minimum cohort sizes to detect changes. Additionally, external QC samples are useful in situations where samples of the same matrix or the same type of sample preparation are used regularly in a laboratory. For example, if total plasma isolated with a K_2_-EDTA anticoagulant is regularly considered in different biological settings, it can be helpful to have a large quantity of pooled plasma that has been well-characterized previously on hand as a control. When prepared alongside experimental samples in each new batch or experiment, you can ensure similar performance relative to past projects.

The same internal QCs that were added to the experimental samples should be added to the external QC samples. For example, protein internal QCs can indicate if the external QC sample’s preparation was comparable to others within a batch or across a plate. Peptide internal QCs, such as PRTC, are used to evaluate whether acquisition was consistent.

It is important to note that the use of internal and external QC samples should be separate from signal calibration. The use of a reference material for single point calibration has been described by our group and others^52,84,85^. We have used internal QCs and the external inter-batch and inter-experiment QC samples to evaluate the performance of normalization, signal calibration, or batch correction. However, the controls described here are not replacements for published calibration techniques. It is important that the standards used for QC of the sample preparation and quantitative analysis are not the same samples that are used in the normalization and correction of the data.

In closing, we have described a series of real-world applications and case studies of QC measures spanning the assessment of system suitability, sample processing, and data analysis. These data have been used to develop recommendations for QA and QC techniques for LC-MS proteomics experiments. Moving beyond identifications and relying on a more integrated, adaptable approach to QC in quantitative proteomics can jumpstart troubleshooting LC-MS systems and can increase confidence in the results of quantitative results.

## Supporting information

Supplementary_figures_tables_methods_discussion

## Author contributions

Conceptualizing (K.A.T., G.E.M., C.C.W., A.N.H., M.J.M.), experimental design (K.A.T., G.E.M., J.E.R., R.S.J., D.L.P, J.D.C., S.E.S.), sample preparation (G.E.M., J.E.R. R.S.J.), data collection (K.A.T., G.E.M., J.E.R., R.S.J., J.D.C., E.H.), data analysis (K.A.T., G.E.M., J.E.R., R.S.J., J.P., J.D.C.), figure generation (K.A.T., J.P.), software development (M.R., V.S., B.X.M, J.E., M.S.B.), writing: original draft (K.A.T., G.E.M.), writing: review and editing (all authors).

## Acknowledgements

We acknowledge the input from current and former MacCoss lab members over the last 20 years who contributed to the evolution of these methods. We also thank Dr. Brook Nunn and her group for valuable feedback on these methods.

This work was supported in part by National Institutes of Health grants (U19 AG065156, R24 GM141156, RF1 AG053959, P30 AG013280, F31 AG069420, and T32 AG066574), and the University of Washington’s Proteomics Resource (UWPR95794).

This research was also supported by the Intelligence Advanced Research Projects Activity (IARPA) TEI-REX program through the Army Research Office contract W911NF2220059. The views and conclusions contained should not be interpreted as necessarily representing the official policies, either expressed or implied, of ODNI, IARPA, ARO, or the U.S. Government. The U.S. Government is authorized to reproduce and distribute preprints for governmental purposes notwithstanding any copyright annotation therein.

## Abbreviations

BSA: bovine serum albumin
CSF: cerebrospinal fluid
CV: coefficient of variance
DDA: data-dependent acquisition
DIA: data-independent acquisition
DTT: dithiothreitol
ENO: yeast enolase
HCD: high energy collisional dissociation
HPLC: high performance liquid chromatography
IAA: iodoacetamide
LC-MS: liquid chromatography mass spectrometry
MCX: mixed-mode ion exchange
PRTC: peptide retention time calibration mixture
PRM: parallel reaction monitoring
PSM: peptide-spectrum match
PTM: post-translational modification
QA: quality assurance
QC: quality control
TEAB: triethylammonium bicarbonate
UPLC: ultra-performance liquid chromatography

## Supporting Information

The supporting material includes additional figures, tables, extended methods descriptions, and a Frequently Asked Questions section.

- **Figure S1:** Using protein and peptide internal QCs to identify a sample preparation issue.
- **Figure S2:** Using system suitability runs and sample peptide internal QCs to identify a system failure.
- **Table S1:** Description and composition of QCs described in framework.
- **Table S2:** Target mass list for PRM system suitability method.
- **Supplementary methods:** In-depth methods description related to the 6 summarized real-world examples.
- **Supplementary discussion and FAQ:** Additional detailed discussion on QC framework development and frequently asked questions.

