## Supplementary_figures_tables_methods_discussion for "A framework for quality control in quantitative proteomics"


### Table of Contents


**Content** **Pages**

**Supplementary Figures and Tables…………………………………………………………… 2-5**

- Figure S1 2
- Figure S2 3
- Table S1 4
- Table S2 5

**Supplementary Methods……………………………………………………………………… 6-12**

**Supplementary Discussion and Frequently Asked Questions………………………………... 13-20**

### Supplementary Figures

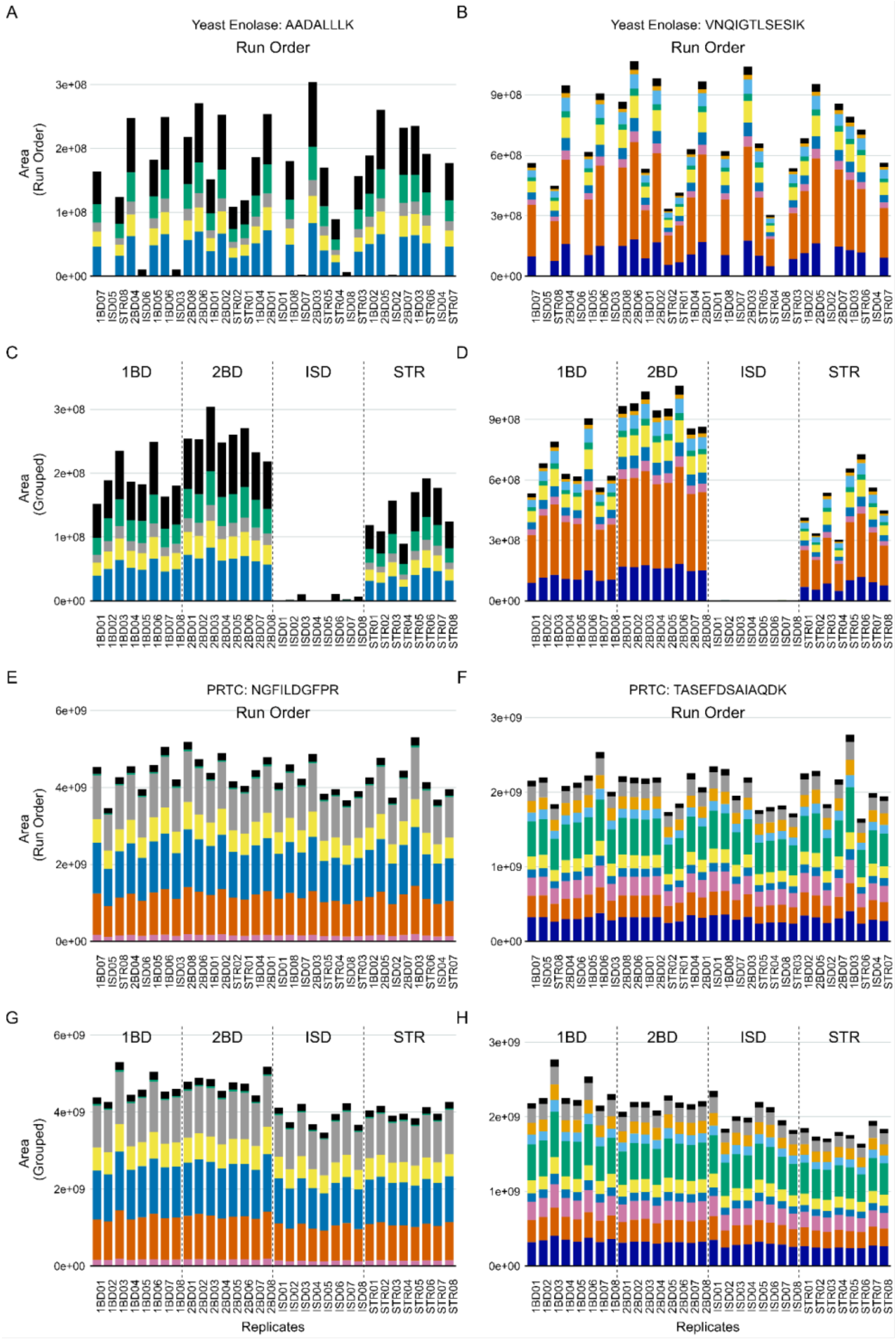


**Figure S1:** The area of two ENO peptides (AADALLLK, VNQIGTLSESIK) and two PRTC peptides (NGFILDGFPR, TASEFDSAIAQDK) representative of observed trends described in Figure 4 are shown here. Peptide VNQIGTLSESIK in panel B and D was also plotted in Figure 4A and 4B. All four peptides are plotted in run order (A, B, E, F) and again grouped by digestion condition (C, D, G, H). ENO peptides were missing in numerous runs (A and B) and when reordered belonged to the ISD sample preparation (C and D). PRTC peptides were broadly consistent across all sample conditions (B, D, F, H). We later found that our RapiGest SF stock, which was one of the only reagents different between the four protocols, was expired and likely impaired the digestion protocol, and thus the ENO, without any impact to the PRTC levels.

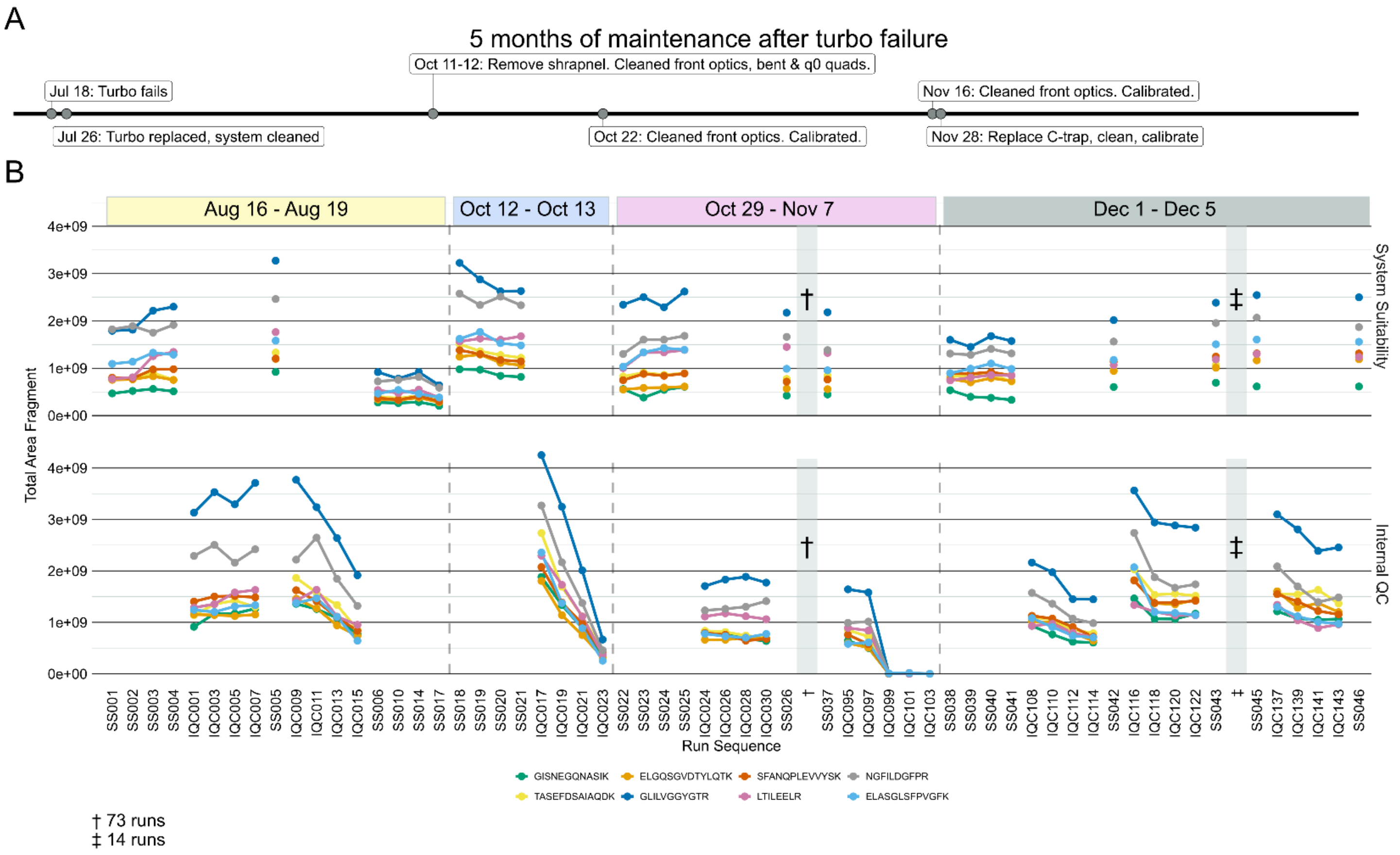


**Figure S2:** Combining system suitability injections and sample internal QCs facilitate real-time monitoring of the system. On an Orbitrap Lumos Tribrid, a significant turbo failure led to months of inconsistent and unpredictable data collection. A rough summary of the major maintenance steps during 5 months of troubleshooting post-turbo failure are shown here (A). Between August - December, signal intensity would unpredictably begin decreasing with each injection. Eight PRTC peptides from system suitability runs (B, top) and sample internal QCs (B, bottom) illustrate this in four different experimental batches. Only a subset of all runs are shown here to improve visibility, but the trends are representative. The full dataset including all runs for PRTC, ENO, and BSA are available on Panorama. In August, after 7 sample injections (IQC001-IQC007) and 5 system suitability injections (SS001-SS005) the system was found to be stable. As additional samples were run (IQC009-IQC015), the signal intensity dropped significantly. This reduced performance was confirmed with system suitability injections (SS006-SS017). The system was taken offline. Additional metal debris from the turbo failure that was missed in earlier maintenance was removed, and the front optics and quads were cleaned. After calibration, the same issue of rapidly declining signal was observed in mid-October (SS018-IQC023). After taking the system down and cleaning the optics again, the system seemed stable for 75 runs (SS026-SS037) until the signal intensity declined rapidly while running samples IQC097-IQC103. The system was taken down again. In late November metal debris was found lodged in the C-trap, and it was replaced along with another thorough cleaning. We then observed a return to expected signal stability and intensity (SS038 through SS046) relative to matched sample batches run prior to the system failing.

**Supplementary Table 1:** Description and composition of quality control steps listed in Figure 1 and applied in the experimental examples encompassing Figures 2-7 and Supplementary Figure 1 and 2.

| QC Step | Description and example composition* | Workflow step tracked |
|---|---|---|
| System suitability | Dedicated LC-MS injections of a standardized sample used for nearly real-time data assessment at the beginning, middle, and end of run sequence. They are also used for longitudinal system tracking in conjunction with digital notebooks, Skyline, and Panorama AutoQC.<br>**Example:**<br>• **Matrix:** 600 fmol BSA peptides/injection<br>• **Peptide internal QC:** 150 fmol PRTC peptides/injection<br>• **PRM acquisition** | System |
| Inter-experiment quality control sample | External quality control sample to assess computational and statistical strategies after adjusting for run order, batch effects, calibration, etc. Prepared alongside each batch of samples. Similar composition, but different pool than the Inter-batch quality control. Can be prepared from analytical samples, or from samples of the same or similar matrix.<br>**Example:**<br>• **Matrix:** Pooled quantity of tissues<br>• **Protein internal QC:** 800 ng yeast enolase/sample<br>• **Peptide internal QC:** 150 fmol PRTC peptides/injection | Quantitative analysis |
| Inter-batch quality control sample | External quality control sample that is prepared alongside each batch of samples to correct for batch-to-batch sample preparation and data collection differences in intensity. Similar composition, but different pool than the Inter-experiment quality control. Generated from samples in an individual sample batch.<br>**Example:**<br>• **Matrix:** Pool of equal quantities of samples<br>• **Protein internal QC:** 800 ng yeast enolase/sample<br>• **Peptide internal QC:** 150 fmol PRTC peptides/injection | Process and Quantitative analysis |
| Peptide internal quality control | Internal quality control. Peptides spiked into samples to monitor system function. When used in conjunction with protein internal quality controls, they can be used to parse out whether the process is the root of an identified issue.<br>**Example:** 50 fmol/µL PRTC peptides. | Process and System |
| Protein internal quality control | Internal quality control. Protein spiked into sample to evaluate sample processing and peptide recovery. May examine digestion or PTM enrichment depending on the protein modifications and stage of introduction. Used in tandem with a peptide internal control, they may also be used to delineate between system issues and sample preparation problems.<br>**Example (digestion):** 800 ng yeast enolase/sample<br>**Example (phosphopeptide enrichment):** 400 ng beta-casein/sample | Process and System |

**Supplementary Table 2:** Target mass list for the parallel reaction monitoring (PRM) system suitability method described and employed in this quality control framework. Includes 15 PRTC peptides and two peptides originating from bovine serum albumin (BSA). All 17 targets were z = 2.

| Peptide | *m/z* | Source | Label |
|---|---|---|---|
| ATEEQLK | 409.7 | In-house digested BSA | light |
| IGDYAGIK | 422.7 | PRTC peptide | heavy |
| DIPVPKPK | 451.2 | PRTC peptide | heavy |
| SSAAPPPPPR | 493.7 | PRTC peptide | heavy |
| HVLTSIGEK | 496.2 | PRTC peptide | heavy |
| LTILEELR | 498.8 | PRTC peptide | heavy |
| GLILVGGYGTR | 558.3 | PRTC peptide | heavy |
| NGFILDGFPR | 573.3 | PRTC peptide | heavy |
| LVNELTEFAK | 582.3 | In-house digested BSA | light |
| SAAGAFGPELSR | 586.8 | PRTC peptide | heavy |
| GISNEGQNASIK | 613.3 | PRTC peptide | heavy |
| ELASGLSFPVGFK | 680.3 | PRTC peptide | heavy |
| TASEFDSAIAQDK | 695.8 | PRTC peptide | heavy |
| SFANQPLEVVYSK | 745.3 | PRTC peptide | heavy |
| ELGQSGVDTYLQTK | 773.8 | PRTC peptide | heavy |
| LSSEAPALFQFDLK | 787.4 | PRTC peptide | heavy |
| GILFVGSGVSGGEEGAR | 801.4 | PRTC peptide | heavy |

### Supplementary Methods

#### Quality Control Sample Composition and Associated Reagents

Figure 1 illustrates the various QCs used (A) and the stages they were introduced (B). For our system suitability control, we typically injected 30-150 fmol Pierce™ Retention Time Calibration (PRTC) Mixture (Thermo Scientific) in a carrier background of 600 fmol of a bovine serum albumin (BSA) tryptic digest in 0.1% trifluoroacetic acid in water. For internal QCs, we typically began by adding 16 ng yeast enolase per µg total protein in the sample (based on a BCA protein assay), which captured variation in the sample processing including tryptic digestion and LC-MS. A second internal QC (30-150 fmol per LC-MS run) of PRTC was added just prior to LC-MS analysis, which captured variation from the LC-MS steps. The quantity of peptide internal QCs were kept consistent between the system suitability and experimental samples. We used two forms of external QC samples: one to assess sample preparation, and one for normalization or batch correction. The external QC samples were composed of two of the following options, with the best options listed first: a pool derived from representative experimental samples, additional samples from the same source as the experimental samples, or the same type of sample but from another source. The same protein and peptide internal QCs were included in the external QC samples.

#### Vignette 1: System Suitability on Orbitrap Eclipse Tribrid

The system suitability standard was loaded onto Kasil1 (PQ Corporation) fritted microcapillary trap (2-3 cm x 150 µm) loaded with 3 µm Reprosil-Pur C18 (Dr. Maisch) reverse-phase beads. Once the sample was loaded to the trap, it was brought in-line with a 30 cm x 75 µm packed tip (New Objective) containing the same resin used for the trap. The high-performance liquid chromatography (HPLC) was a Thermo Easy nLC1200 and used 0.1% formic acid in water as buffer A, and 0.1% formic acid /80% acetonitrile/19.9% water as buffer B. The strong needle wash was 50% acetonitrile/50% water, and the weak needle was 0.1% trifluoroacetic acid/2% acetonitrile/97.9% water. The 50-minute system suitability gradient consisted of 2 to 40% B in 30 minutes, 40 to 75% in 5 minutes, 75 to 100% B in 15 minutes, followed by a wash and a 30-minute column equilibration. The trap and column were maintained at a constant temperature of 50°C within a heated source (CorSolutions) and electrosprayed into a Thermo Orbitrap Eclipse Tribrid Mass Spectrometer with the application of a distal 2 kV spray voltage. The scan cycle included one 30,000 resolution full-scan mass spectrum (*m/z* 400-810) followed by 17 PRM MS/MS spectra targeting 2 BSA peptides and the 15 heavy-labeled PRTC peptides at 15,000 resolution, AGC target of 5e4, 22 ms maximum injection time, and 30% normalized collision energy with a *m/z* 2 isolation window.

The target mass list for the system suitability method included 17 entries (all z = 2) that are listed in Supplementary Table 2.

#### Vignette 2: MS issue on Orbitrap Fusion

The PRTC/BSA system suitability standard was as described above. The tryptic yeast proteome was obtained from Promega, which was reduced, alkylated, and digested with trypsin according to the manufacturer's instructions.

For the yeast DDA runs (n = 3 technical replicates per batch), one µg of yeast peptides were loaded, and for the PRM system suitability runs (5-11 technical replicates per batch), 3 µl of our system suitability standard was injected.

For both the DDA and PRM methods, a Waters NanoAcquity UPLC was coupled to a Thermo Orbitrap Fusion. Mobile phase buffer A was 0.1% formic acid in water and buffer B was 0.1% formic acid in 100% acetonitrile. The strong needle wash was 50% acetonitrile/50% water, and the weak needle was 0.1% trifluoroacetic acid/2% acetonitrile/97.9% water. For DDA runs, the 90-95 minute gradient began with 2% B and consisted of 2 to 7.5% B in 10 minutes, 7.5 to 25% in 40 minutes, 25 to 60% B in 25 minutes, 60 to 95% B in 6 minutes, followed by column equilibration with 2% B for 14 minutes. For PRM runs, the 55-60 minute gradient began with 2% B and consisted of 2 to 35% B in 30 minutes, 35 to 60% in 11 minutes, 60 to 95% B in 6 minutes, followed by column equilibration with 2% B in 13 minutes. The trap and column were maintained at a temperature of 50°C using a heated source (CorSolutions) and electrosprayed into a Thermo Orbitrap Fusion Mass Spectrometer with the application of a distal 2 kV spray voltage.

For the DDA runs, a cycle of one 120,000 resolution MS1 scan (*m/z* 400-1600) was followed by data-dependent high energy collisional dissociation (HCD) fragmentation to acquire MS/MS spectra targeting the most intense ions. In the precursor scan, we used a charge state filter of +2 to +7, and sampled peaks were added to a dynamic exclusion list with a 20 second duration, the AGC target was 200,000, and a 20 ms maximum injection time. The MS/MS spectra (*m/z* 100-1000) acquired in the linear ion trap used an AGC target of 10,000, a 35 ms maximum injection time, and 30% normalized collision energy. For the PRM runs, a precursor scan (*m/z* 400-1600) was followed-up with a set of targeted MS/MS scans. MS/MS spectra were acquired in a linear ion trap using low energy beam CID (HCD) fragmentation, with an AGC target of 10,000, a maximum injection time of 35 s, and a 30% normalized collision energy.

We assessed the chromatography, retention times, and peak areas of 15 PRTC peptides and 2 BSA peptides in the PRM system suitability runs using Skyline (version 22.2). The raw data, Skyline document, Skyline report files, and the processed data used as input to generate Figure 3 are available on PanoramaWeb.

Using a Nextflow pipeline (https://github.com/mriffle/nf-teirex-dda, revision 068f68323c9f9a181175a81ee796ef4a3373b5ed), the DDA MS data were converted to mzML with msConvert[57], peptides identified using Comet[58,59], version 2023.01 rev. 2 (uwpr.github.io/Comet/releases/release_202301), q-values and posterior error probabilities at the peptide-spectrum match (PSM) level were acquired using Percolator[60] version 3.06 (github.com/percolator/percolator/releases), and uploaded to Limelight[61] for visualization and dissemination. Searches were performed using a Saccharomyces cerevisiae reference proteome FASTA file (Uniprot Proteome ID: UP000002311, downloaded January 27, 2024) appended with the internal control yeast enolase 1 and a common list of contaminants generated in-house. The raw data, parameter files used in Nextflow DDA analyses, and the processed data downloaded from Limelight used as input to generate Figure 3 are available on PanoramaWeb. The analyzed DDA data is freely accessible on Limelight (Project ID = 131).

### Vignette 3: Sample processing issue on Orbitrap Lumos Tribrid

Sample preparations of commercial human CSF (Golden West Biologicals) included four different approaches. Two were variations on the paramagnetic bead-based, single-pot solid-phase-enhanced sample preparation (SP3) that was developed and optimized by others[62–65] (labels: 1BD and 2BD). The other two methods tested included S-trap column digestion and clean-up (label: STR), and in-solution digestion with Rapigest SF (label: ISD), and mixed-mode solid phase extraction clean-up (label: MIX). A large pool of Golden West human CSF was aliquoted into 50 µL volumes of 8 replicates for each sample preparation type.

For the 1BD SP3 sample preparation each 50 µL aliquot of human CSF was resuspended in 8M Urea, 150 mM NaCl, 100 mM Tris pH 8.5 with 800 ng of yeast enolase protein (Sigma-Aldrich) added as an internal protein control. Processing included reduction with 10 mM dithiothreitol (DTT) (30°C, 10 minutes) and alkylation with 15 mM iodoacetamide (IAA) (25°C, 30 minutes). Proteins were aggregated on 50 µL of 10 µg/µL MagReSyn carboxylate beads (ReSyn Biosciences) with 50% ethanol/50% water, washed three times with 80% ethanol/20% water, and digested to peptides with trypsin (1:25) in 25 mM ammonium bicarbonate, using a Thermo KingFisher Flex. Samples were acidified to a final concentration of 0.1% formic acid in water.

For the 2BD SP3 sample preparation each 50 µL aliquot of human CSF was resuspended in 2% SDS, 100 mM Tris pH 8.5 with 800 ng of yeast enolase protein (Sigma-Aldrich) added as an internal protein control. Processing included reduction with 10 mM DTT (60°C, 10 minutes) and alkylation with 15 mM IAA (25°C, 30 minutes). Proteins were aggregated on 25 µL of 20 µg/µL MagReSyn hydroxyl beads (ReSyn Biosciences) with 70% acetonitrile, washed three times with 95% acetonitrile/5% water, washed twice with 70% ethanol/30% water, and digested to peptides with trypsin (1:25) in 50 mM ammonium bicarbonate, using a Thermo KingFisher Flex. Samples were acidified to a final concentration of 0.1% formic acid in water.

For the STR sample preparation each 50 µL aliquot of human CSF was resuspended in 50 µL of 2X lysis buffer of 10% SDS, 100 mM triethylammonium bicarbonate (TEAB) with 2X Halt protease and phosphatase inhibitors (Thermo Scientific) with 800 ng of yeast enolase protein (Sigma-Aldrich) added as an internal protein quality control. Processing included reduction with 20 mM DTT (60°C, 10 minutes) and alkylation with 40 mM IAA (25°C, 30 minutes). Lysates were prepared for S-trap column (Protifi) binding by the addition of 1.2% phosphoric acid and 350 µL of binding buffer (90% methanol, 100 mM TEAB). The acidified lysate was bound to the column incrementally. This was followed by 3 wash steps with binding buffer to remove SDS, 3 wash steps with 50:50 methanol:chloroform to remove lipids, and a final wash step with binding buffer. Trypsin (1:25) in 50mM TEAB was added to the S-trap column for digestion at 47°C for one hour. Hydrophilic peptides were eluted with 50 mM TEAB followed by hydrophobic peptides with a solution of 0.2% formic acid/50% acetonitrile/49.8% water. Eluates were pooled, speed vacuumed, and resuspended in 0.1% formic acid in water.

For the ISD sample preparation each 50 µL aliquot of human CSF was resuspended in 0.2% RapiGest SF (Waters) in 50 mM ammonium bicarbonate with 800 ng of yeast enolase protein (Sigma-Aldrich) added as an internal protein control. Processing included reduction with 10 mM DTT (60°C, 30 minutes), alkylation with 15 mM IAA (25°C, 30 minutes) and quenching with 10 mM DTT. Samples were digested in trypsin (1:25) in 50 mM ammonium bicarbonate at 37°C for 16 hours. Samples were quenched with 200 mM HCl and cleaned with mixed-mode ion exchange (MCX) columns (Waters).

One µg of each digested sample and 150 femtomole of Pierce Retention Time Calibrant (PRTC) was loaded onto a 150 µm fused silica Kasil1 (PQ Corporation) fritted microcapillary trap loaded with 3.5 cm of 3 µm Reprosil-Pur C18 (Dr. Maisch) reverse-phase resin coupled with an 75 µm inner diameter picofrit (New Objective) analytical column containing 30 cm of 3 µm Reprosil-Pur C18 attached to a Thermo EASY-nLC 1200. The PRTC was also used to assess the quality of the column before and during analysis. We analyzed four of these system suitability runs prior to any sample analysis. After every six to eight sample runs, another system suitability run was analyzed. Buffer A was 0.1% formic acid in water and buffer B was 0.1% formic acid/80% acetonitrile/19.9% water. The strong needle wash was 50% acetonitrile/50% water, and the weak needle was 0.1% trifluoroacetic acid/2%

acetonitrile/97.9% water. The 40-minute system suitability gradient consisted of 0 to 16% B in 5 minutes, 16 to 35% in 20 minutes, 35 to 75% B in 5 minutes, 75 to 100% B in 5 minutes, followed by a wash of 9 minutes and a 30-minute column equilibration. The 110-minute sample LC gradient consists of a 2 to 7% for 1 minute, 7 to 14% B in 35 minutes, 14 to 40% B in 55 minutes, 40 to 60% B in 5 minutes, 60 to 98% B in 5 minutes, followed by a 9-minute wash and a 30-minute column equilibration. Peptides were eluted from the column with a 50°C heated source (CorSolutions) and electrosprayed into a Thermo Orbitrap Fusion Lumos Mass Spectrometer with the application of a distal 3 kV spray voltage. For the system suitability analysis, a cycle of one 120,000 resolution full-scan mass spectrum (350-2000) followed by a data-independent MS/MS spectra on the loop count of 76 data-independent MS/MS spectra using an inclusion list at 15,000 resolution, AGC target of 4e5, 20 millisecond (ms) maximum injection time, 33% normalized collision energy with an *m/z* 8 isolation window. For the sample digest, data were collected using data-independent acquisition (DIA) strategies. First, a chromatogram library of 6 independent injections was collected from a pool of all samples within a batch. For each injection, a cycle of one 120,000 resolution full-scan mass spectrum with a mass range of *m/z* 110 (*m/z* 395-505, *m/z* 495-605, *m/z* 595-705, *m/z* 695-805, *m/z* 795-905, or *m/z* 895-1005) followed by a data-independent MS/MS spectra on the loop count of 26 at 30,000 resolution, AGC target of 4e5, 60 ms maximum injection time, 33% normalized collision energy with a *m/z* 4 overlapping isolation window. The chromatogram library data was used to quantify proteins from individual sample runs. These individual runs consisted of a cycle of one 120,000 resolution full-scan mass spectrum with a mass range of *m/z* 350-2000, AGC target of 4e5, 100 ms maximum injection time followed by a data-independent MS/MS spectra on the loop count of 76 at 15,000 resolution, AGC target of 4e5, 20 ms maximum injection time, 33% normalized collision energy with an overlapping *m/z* 8 isolation window. Application of the mass spectrometer and LC solvent gradients were controlled by the ThermoFisher Xcalibur data system.

System suitability and internal controls in samples and external QC samples were imported into Skyline (version 22.2) using similar settings for PRM and DIA data.

#### Vignette 4: LC issue on Orbitrap Eclipse Tribrid

Human plasma was resuspended in 2% SDS, 50 mM Tris pH 8.5 with 1X Halt protease and phosphatase inhibitors (Thermo Scientific). Protein concentration was measured using the Pierce BCA assay (Thermo Scientific). Samples were prepared with 50 µg of homogenate and 800 ng of yeast enolase protein (Sigma) as an internal protein control. Processing included reduction with 10 mM DTT (60°C, 10 minutes) and alkylation with 15 mM IAA (25°C, 30 minutes). Proteins were aggregated on MagReSyn Hydroxyl beads (ReSyn Biosciences), washed (3 times with 95% acetonitrile/5% water, and 2 times with 70% ethanol/30% water), and digested to peptides, in a 96-well plate using a Thermo KingFisher Flex. Samples were acidified to a final concentration of 0.1% formic acid in water.

One µg of each digested sample and 150 femtomole of Pierce Retention Time Calibrant (PRTC) was loaded onto a 150 µm fused silica Kasil1 (PQ Corporation) fritted microcapillary trap loaded with 3.5 cm of 3 µm Reprosil-Pur C18 (Dr. Maisch) reverse-phase resin coupled with an 75 µm inner diameter picofrit (New Objective) analytical column containing 30 cm of 3 µm Reprosil-Pur C18 attached to a Thermo EASY-nLC 1200. The PRTC was also used to assess the quality of the column before and during analysis. We analyzed four of these system suitability runs prior to any sample analysis. After every eight sample runs, another system suitability run was analyzed. Buffer A was 0.1% formic acid in water and buffer B was 0.1% formic acid/80% acetonitrile/19.9% water. The strong needle wash was 50% acetonitrile/50% water, and the weak needle was 0.1% trifluoroacetic acid/2%

acetonitrile/97.9% water. The 50-minute system suitability gradient consisted of 2 to 40% B in 30 minutes, 40 to 75% in 5 minutes, 75 to 100% B in 15 minutes, followed by a wash and a 30-minute column equilibration. The 100-minute sample LC gradient consists of a 2 to 40% for 80 minutes, 40 to 75% B in 10 minutes, 75 to 100% B in 10 minutes, followed by a 30-minute wash and column equilibration. Peptides were eluted from the column with a 50°C heated source (CorSolutions) and electrosprayed into a Thermo Eclipse Tribrid Mass Spectrometer with the application of a distal 3 kV spray voltage. For the system suitability analysis, a cycle of one 30,000 resolution full-scan mass spectrum (*m/z* 400-810) followed by a data-independent MS/MS spectra on the loop count of 20 data-independent MS/MS spectra using an inclusion list at 15,000 resolution, AGC target of 5e4, 22 ms maximum injection time, 30% normalized collision energy with a *m/z* 2 isolation window with a mass list table for the 2 BSA peptides and the 15 heavy labeled PRTC peptides. For the sample digest, first a chromatogram library of 6 independent injections was analyzed from a pool of all samples within a batch. For each injection, a cycle of one 30,000 resolution full-scan mass spectrum with a mass range of *m/z* 110 (*m/z* 395-505, *m/z* 495-605, 595- *m/z* 705, *m/z* 695-805, *m/z* 795-905, or *m/z* 895-1005) followed by a data-independent MS/MS spectra on the loop count of 25 at 30,000 resolution, AGC target of 5e5, 54 ms maximum injection time, 27% normalized collision energy with a *m/z* 4 overlapping isolation window. The chromatogram library data was used to quantify proteins from individual sample runs. These individual runs consisted of a cycle of one 30,000 resolution full-scan mass spectrum with a mass range of *m/z* 350-1005, AGC target of 4e5, 50 ms maximum injection time followed by a data-independent MS/MS spectra on the loop count of 75 at 15,000 resolution, AGC target of 4e5, 22 ms maximum injection time, 27% normalized collision energy with an overlapping *m/z* 8 isolation window. Application of the mass spectrometer and LC solvent gradients were controlled by the ThermoFisher Xcalibur data system.

System suitability and internal controls in samples and external QC samples were imported into Skyline (version 22.2) using similar settings for PRM and DIA data.

#### Vignette 5: Combining system suitability and internal quality controls on Orbitrap Lumos Tribrid

Mouse brain micropunches were homogenized in 30 µL of 5% SDS, 50 mM triethylammonium bicarbonate (TEAB) with 1X Halt protease and phosphatase inhibitors (Thermo Scientific) in a Barocycler 2320 EXT (Pressure Biosciences Inc.) for 30 repeat cycles (20 sec at 45k psi, 10 sec at ambient pressure) at 35°C. Protein concentration was measured using the Pierce BCA assay (Thermo Scientific). Samples were prepared with 25 µg of homogenate and 400 ng of yeast enolase protein (Sigma) as an internal protein control. Processing included reduction with 20 mM DTT (60°C, 10 minutes) and alkylation with 40 mM IAA (25°C, 30 minutes). Lysates were prepared for S-trap column (Protifi) binding by the addition of 1.2% phosphoric acid and 350 µL of binding buffer (90% methanol, 100 mM TEAB). The acidified lysate was bound to the column incrementally. This was followed by 3 wash steps with binding buffer to remove SDS, 3 wash steps with 50:50 methanol:chloroform to remove lipids, and a final wash step with binding buffer. Trypsin (1:10) in 50mM TEAB was added to the S-trap column for digestion at 47°C for one hour. Hydrophilic peptides were eluted with 50 mM TEAB followed by hydrophobic peptides with a solution of 0.2% formic acid/50% acetonitrile/49.8% water. Eluates were pooled, speed vacuumed, and resuspended in 0.1% formic acid in water.

One µg of each digested sample and 150 femtomole of Pierce Retention Time Calibrant (PRTC) was loaded onto a 150 µm fused silica Kasil1 (PQ Corporation) fritted microcapillary trap loaded with 3.5 cm of 3 µm Reprosil-Pur C18 (Dr. Maisch) reverse-phase resin coupled with an 75 µm inner diameter picofrit (New Objective) analytical column containing 30 cm of 3 µm Reprosil-Pur C18 attached to a

Thermo EASY-nLC 1200. The PRTC was also used to assess the quality of the column before and during analysis. We analyzed four of these system suitability runs prior to any sample analysis. After every six to eight sample runs, another system suitability run was analyzed. Buffer A was 0.1% formic acid in water and buffer B was 0.1% formic acid/80% acetonitrile/ 19.9% water. The strong needle wash was 50% acetonitrile/50% water, and the weak needle was 0.1% trifluoroacetic acid/2% acetonitrile/97.9% water. The 40-minute system suitability gradient consisted of 0 to 16% B in 5 minutes, 16 to 35% in 20 minutes, 35 to 75% B in 5 minutes, 75 to 100% B in 5 minutes, followed by a wash of 9 minutes and a 30-minute column equilibration. The 110-minute sample LC gradient consists of a 2 to 7% for 1 minute, 7 to 14% B in 35 minutes, 14 to 40% B in 55 minutes, 40 to 60% B in 5 minutes, 60 to 98% B in 5 minutes, followed by a 9-minute wash and a 30-minute column equilibration. Peptides were eluted from the column with a 50°C heated source (CorSolutions) and electrosprayed into a Thermo Orbitrap Fusion Lumos Mass Spectrometer with the application of a distal 3 kV spray voltage. For the system suitability analysis, a cycle of one 120,000 resolution full-scan mass spectrum (*m/z* 350-2000) followed by a data-independent MS/MS spectra on the loop count of 76 data-independent MS/MS spectra using an inclusion list at 15,000 resolution, AGC target of 4e5, 20 millisecond (ms) maximum injection time, 33% normalized collision energy with an *m/z* 8 isolation window. For the sample digest, first a chromatogram library of 6 independent injections was collected from a pool of all samples within a batch. For each injection, a cycle of one 120,000 resolution full-scan mass spectrum with a mass range of *m/z* 110 (*m/z* 395-505, *m/z* 495-605, *m/z* 595-705, *m/z* 695-805, *m/z* 795-905, or *m/z* 895-1005) followed by a data-independent MS/MS spectra on the loop count of 26 at 30,000 resolution, AGC target of 4e5, 60 ms maximum injection time, 33% normalized collision energy with a *m/z* 4 overlapping isolation window. The chromatogram library data was used to quantify proteins from individual sample runs. These individual runs consisted of a cycle of one 120,000 resolution full-scan mass spectrum with a mass range of *m/z* 350-2000, AGC target of 4e5, 100 ms maximum injection time followed by a data-independent MS/MS spectra on the loop count of 76 at 15,000 resolution, AGC target of 4e5, 20 ms maximum injection time, 33% normalized collision energy with an overlapping *m/z* 8 isolation window. Application of the mass spectrometer and LC solvent gradients were controlled by the ThermoFisher Xcalibur data system.

System suitability and internal controls in samples and external QC samples were imported into Skyline (version 22.2) using similar settings for PRM and DIA data.

#### Vignette 6: Assessing quantitative results on Orbitrap Eclipse Tribrid with external quality controls

Lumbar CSF from 280 patients were divided into four major groups: 1) Healthy Control, 2) Alzheimer's disease/mild cognitive impairment, 3) Parkinson's disease cognitively normal and 4) Parkinson's disease cognitively impaired. Each row of half a 96-well plate contained 10 balanced and randomized CSF samples and two external QC samples. One external inter-batch QC sample was a pool of 50 patients representing all 4 groups and the other external inter-experiment QC sample was a commercially available pool of CSF. These controls were processed with the samples and used to evaluate the technical precision within and between each batch prior to and following normalization and batch adjustment.

The CSF samples (25 µg) were resuspended in SDS lysis buffer (working concentration of 1% SDS, 50 mM Tris, pH 8.5) with 400 ng of yeast enolase as a protein internal control to assess sample digestion. Experimental samples and external QC samples were reduced with 40 mM DTT (10 minutes at 60 °C), alkylated with 80 mM IAA (30 minutes at room temperature), and quenched with 40 mM DTT. Proteins were aggregated on MagReSyn Hydroxyl beads by diluting samples to 70%

acetonitrile/30% water, washed three times with 95% acetonitrile/5% water, washed twice with 70% ethanol/30% water, and digested to peptides with 1:10 trypsin at 47°C for one hour using a Thermo KingFisher Flex. Samples were spiked with 150 fmol of PRTC per injection volume. Peptides were separated using reverse-phase chromatography with a Thermo Easy nano-LC and electrosprayed into a Thermo Orbitrap Eclipse Tribrid analyzed using a *m/z* 12 staggered DIA isolation scheme. The data were demultiplexed to *m/z* 6 with ProteoWizard with settings of "overlap_only" and Mass Error = 10.0 ppm. A Prosit library is generated using the Uniprot human canonical FASTA (Proteome ID: UP000005640) and NCE setting=33[66,67]. Using EncyclopeDIA (version 2.12.30), the 6x GPF-DIA acquisitions from each plate batch are combined to a single global GPF library, shifts in retention times are aligned, and batch-specific GPF libraries are generated. The Prosit library is empirically corrected by searching the batch-specific GPF libraries as described previously[68,69]. Default EncyclopeDIA settings are used: (10 ppm tolerances, trypsin digestion, and higher-energy collisional dissociation [HCD] b- and y-ions). The results from this analysis were saved as a "Chromatogram Library" in EncyclopeDIA's eLib format. The "wide-window" DIA runs are analyzed using EncyclopeDIA requiring a minimum of 3 quantitative ions and filtering peptides with q-value ≤ 0.01 using Percolator 3.01. After analyzing each file individually, EncyclopeDIA is used to generate a "Quant Report" which stores all the detected peptides, integration boundaries, quantitative transitions, and statistical metrics from all runs in an eLib format.

The Quant Report eLib library was imported into Skyline using the human Uniprot FASTA as the background proteome to map peptides to proteins, perform peak integration, manual evaluation, and generate reports. The sample mzML files were then imported into Skyline. A csv file of peptide level total area fragments (TAFs) for each replicate was exported from Skyline using the custom reporting capabilities of the document grid. Peptide quantities were then median normalized, and batch corrected. All levels of data are available via PanoramaWeb. The coefficient of variation (CV) calculated for all peptides in the inter-batch QC improved 6.3% after normalization and batch adjustment at the peptide-level and 16.9% at the protein level. We also used internal process controls to assess sample preparation and monitor data collection. These controls were necessary, as the data collected encompassed seven batches analyzed on separate LC columns and traps over a three-month period.

### Data Accessibility and Figure Generation

Raw files (DIA, PRM, DDA), Skyline documents, processed results (DIA, PRM) used as input for figure generation, FASTA files, EncyclopeDIA files, and metadata are available on PanoramaWeb (panoramaweb.org/maccoss-sample-qc-system-suitability.url). The data DOI is 10.6069/hmtb-tp04. The processed DDA results in Figure 3 are freely accessible on Limelight[61] under the identifier Project 131. The dataset are also registered through the ProteomeXchange with the unique identifier PXD051318.

The Inkscape files and Panorama AutoQC images used to generate Figures 1 and 2 are available on Panorama and GitHub. Inkscape was used for minor aesthetic changes, panel labeling, and resizing of figures. Input files for R scripts >25 MB are available on PanoramaWeb. The input files <25 MB and the R code to generate Figures 3-6 (maccoss_qc_figures_3_4_5_6_S1.Rmd) and Figure 7 (plot_7a.r, plot_7b.r) are freely available for download on GitHub: (github.com/uw-maccosslab/manuscript-qc-system-suitability/).

### Supplementary Discussion and Frequently Asked Questions

Here we share additional discussion regarding frequently asked questions about the development and practical application of the QC framework described and its associated controls.

1. **What historical steps and selection criteria did the authors use to optimize, validate, and select the composition of the 3 types of quality control approaches described in the framework?**
    a. **System Suitability**
    **<u>Selection criteria for system suitability sample:</u>**
        - Stable (Sample does not change with time during storage or on autosampler between injections)
            - Evaluate peptide stability over the anticipated time that system suitability sample would be left on autosampler.
            - Compare findings between different reagent lots and plan accordingly to always leave sample in reserve for testing between lots.
        - Moderately complex
            - Similar complexity to samples.
        - Minimal carryover
            - Will not dirty instrument or impact next measurement.
        - Commercially available
            - Selection will be dependent upon the stringency of the vendor's quality control – needs to be verified by the user.
        - Inexpensive
            - Ideally. Our group compromises on using a less expensive protein (BSA) as a matrix to include more costly PRTC peptides that are matched to the quantity injected in sample runs.

    **<u>Selection criteria for system suitability instrument method:</u>**
        - Acquisition method
            - We prefer targeted PRM methods for the following reasons:
                - No additional database searching, which is software is regularly updated or settings are changed, can affect how the system suitability data is interpreted.
                - Better for evaluating peak shape, chromatography, and reproducibility.
                - No missing data if there are no changes to the LC gradient (or issues with a batch of reagent, which would ideally have been benchmarked before using for system suitability)
                - In our hands, identified issues more often and more clearly than an identification-based method with DDA.
        - Continuity among different instrument platforms
            - Same or as close to the same LC and MS method used across systems.
                - For 60-90 minute DIA experimental runs across 5 different systems including orbitrap and linear ion trap mass analyzers, we used a 30 minute PRM system suitability method.

  - For 30 minute DIA experimental runs on an Astral, we have experimented with 10 and 20 minute PRM system suitability methods.

**Development of current system suitability sample and methods:**
Previously, we have tried different instrumentation methods and numerous commercial and in-house sample types to evaluate system suitability. More than a decade ago, we experimented with utilizing frequent DDA-based injections of a commercially-available yeast proteome and of a 6 Bovine Tryptic Digest Equal Molar Mix (Michrom Bioresources). This bovine digest mix was discontinued. We moved to a hybridized approach with two commercial products combined to generate a system suitability matrix that we use on all our LC-MS platforms. We digest bovine serum albumin in-house with trypsin and benchmark between lots (protein quantity, digestion efficiency, peptide signal intensity and reproducibility). PRTC, which is also used as a peptide internal QC in our samples, is added in the same quantity (15-150 fmol depending on the time period). The quantity of BSA and PRTC peptides were determined by injecting different quantities onto a system and selecting quantities that generated reasonably intense signal without overloading the column, but not so low as to lose all signal with only a modest decline in system function.

We showed one example in Figure 3 why we shifted to PRM (targeted) system suitability methods in part because the DDA-based identification metric to evaluate system function would not reliably identify even major system failures as we described. Using targeted assays, we do not have additional database searches beyond extracting the known peptides. We can evaluate peak shape and reproducibility without concern for missing data which may occur in DDA-based approaches.

**b. Internal Quality Control**
We use similar selection criteria for internal QCs at the protein and peptide-level.

**Selection criteria for protein and peptide internal QC:**

- Stable
  - Peptides do not change with time during storage or on autosampler between injections.
- Number of peptides used or generated as a result of tryptic digestion (>6)
- Amphipathic peptide coverage of LC-MS gradient (Digested peptides covered the early, middle, and later stages of the LC gradient)
- Reproducibility
  - **Protein-level:** Protein is carried through the experimental protocol every time, digested completely (No missed cleavages), and reliably produces the same peptides across replicates.
  - **Peptide-level:** The same quantity of introduced peptides produces the same signal across replicates of the same matrix.
- Peptides produce strong signal and reproducible peak shape.
- Peptide sequences are not shared with analytically crucial endogenous peptides from the species you are working with.
  - **Protein-level:** Suggest identifying commercially-available protein meeting above qualifications from disparate species.
  - **Peptide-level:** Suggest non-naturally occurring synthetic peptide.

- Indexed retention time (iRT)
    - Do you anticipate wanting or needing to use iRT?

**i. Protein**

**<u>Development of current protein internal QC:</u>**

To evaluate protein digestion, we initially used a $^{15}N$-labeled Apolipoprotein A1 ($^{15}N$-ApoA1) as described in the citation below:

> Henderson, C. M.; Vaisar, T.; Hoofnagle, A. N. Isolating and Quantifying Plasma HDL Proteins by Sequential Density Gradient Ultracentrifugation and Targeted Proteomics BT - Quantitative Proteomics by Mass Spectrometry; Sechi, S., Ed.; Springer New York: New York, NY, 2016; pp 105–120. https://doi.org/10.1007/978-1-4939-3524-6_7.

The $^{15}N$-ApoA1 protein worked well for us in terms of the above selection criteria that we evaluated. However, the $^{15}N$-ApoA1 was costly to use at scale. Eventually, the $^{15}N$-ApoA1 was no longer commercially available.

We sought a more affordable, less likely to be discontinued option. Two yeast proteins were evaluated in terms of the above metrics used to select $^{15}N$-ApoA1: enolase 1 and alcohol dehydrogenase from *Saccharomyces cerevisiae*. We found that both proteins performed well in terms of the quantity of peptides produced after digestion, amphipathic peptide coverage of LC-MS gradient, and peptide reproducibility. At the time we conducted these tests, the protein cost was similar. However, we found that the peptides generated by the yeast alcohol dehydrogenase were more conserved with the mammalian species we frequently work with (humans, mice, companion dogs), and so the group opted to proceed with the yeast enolase instead.

One key drawback to the use of yeast enolase as a protein internal QC is that if you work in *Saccharomyces cerevisiae* or a closely-related organism, an alternative protein will be necessary. In such a scenario, yeast enolase would not produce a reliable, sample processing-related signal due to the contribution of signal from endogenous peptides of the same sequence.

**ii. Peptide**

**<u>Development of current peptide internal QC:</u>**

We have explored the use of labeled synthetic peptides for use as peptide internal QCs, but found that it was challenging for us to internally QC the labeling efficiency between lots. Thus, we moved on to consider two different commercial products with capability to also be used as indexed retention time (iRT) standards:

- Biognosys iRT Kit
    - 11 non-naturally occurring synthetic peptides spanning gradient.
- Pierce Retention Time Calibration Peptides (PRTC)
    - 15 non-naturally occurring synthetic peptides spanning gradient.

Both were costly, but we wanted to move towards a commercial, labeled peptide mixture with quality control more stringent than we could do in-house with regularity. We proceeded with using PRTC due to the higher number of peptides in the mixture, simply because we have found that some peptides do not perform optimally in different matrixes and wanted to maximize our chances of seeing the most internal QC peptides in samples possible.

#### c. External Quality Control Sample

We use similar selection criteria for internal QCs at the protein and peptide-level.

<u>**Selection criteria for inter-batch and inter-experiment external QC samples:**</u>

- Same species as experimental samples (whenever possible)
- Protein concentration within range acceptable for methods
- Minimal freeze-thaws
  - Keep consistent between experiments.
- Reasonable degree of hemolysis for blood products
- Optimally, the same sample type (matrix) as experimental samples
  - For inter-batch QC sample, we suggest pooling experimental sample (If available) or planning to collect additional materials explicitly for use as a control sample.
  - For inter-experiment QC sample, if the same sample matrix is not possible to acquire, strongly suggest a related tissue or biofluid from the same species as your analyte of interest. Two real-world examples are presented – one that worked and one cautionary tale.
    - **Viable alternative:** Experimental samples were mouse hippocampus. We used pooled hippocampus samples for the inter-batch QC sample, but we could not get sufficient tissues to pool for inter-experiment QC sample. Matched mouse cerebellum was available – we found that the preparation and protein results were similar enough and viable for inter-experiment QC sample.
    - **Option that did not work:** When conducting an experiment with lumbar CSF, we considered using a commercial pool of ventricular CSF that we had on hand. We quickly discovered two issues. First - the protein concentration of the commercial pool of ventricular CSF was too low to be useful for the quantity of lumbar plasma we planned to use. Second – the peptides that we identify in lumber and ventricular CSF were so different, it was like using a completely different matrix. We opted not to use this pool as an inter-experiment QC sample for experiments using lumbar CSF.
- For inter-batch QC sample:
  - Representative of sample group.
    - If pooling additional samples rather than portions of experimental samples, ensure that the additional sample is representative of the conditions in your experiment.
    - Example: If you are studying a phenotype in young and aged mice (both sexes), do not only pool young male samples as a quality control. Pool material representative of all groups (1 young male, 1 aged male, 1 young female, 1 aged female), and evenly whenever is feasible.
- For inter-experiment QC sample:
  - Well-characterized and evaluated QC metrics.
    - Previous baseline data collection suggested to better evaluate preparation.
  - Verify that the sample preparation method is compatible with how the pooled inter-experiment QC sample was collected.

- Example: If using Mag-Net to enrich for membrane-bound particles, filtering the plasma drastically decreases the EV yield. Then using a pool of plasma that has been filtered as an external QC sample will not be useful.

**Development of inter-batch and inter-experiment external QC samples:**
Other quantitative analytical methods such as metabolomics can utilize analytical standards to generate a quantity from their MS signal and perform calibration and normalization. In proteomics, especially in discovery-based studies, we cannot utilize standards of all (or often even a few) proteins or peptides of interest due to cost and availability of reagents. Since we cannot reasonably generate calibration curves for the often many thousands of peptides we detect in quantitative proteomics experiments, we effectively substitute the calibration curves that would be used in other approaches with these external QC samples.

We needed a way to evaluate how reliable our sample MS signal was and how well our normalization and batch-correction worked. We postulated that by using repeated measurements of a "known-unknown" sample that is prepared with our experimental samples, we could evaluate these metrics independently of individual sample variation.

These controls are newer for our group and have not been through as many iterations. For the inter-batch control, we use pooled samples that were either collected alongside the experimental samples (whenever feasible) or pooled from small quantities of our experimental samples. This control is used to evaluate variation among and within our sample batches based on peptide and protein-level CVs and correlation among batches. The inter-experiment control is used to evaluate how normalization and batch correction approaches affect overall variation. We expect that when the same sample is processed and analyzed the same way, the variation should be low to begin with, but we would also assume that the variance will not increase after normalization and batch correction (where applicable). This also serves as a general "sanity check" that we have not normalized "too aggressively" and effectively erased any potential for extracting more variable biological phenotypes from the data.

2. **How was AutoQC developed, how do I set-up AutoQC in my laboratory, and what kind of metrics can we evaluate in our samples beyond peak areas, retention times, and mass error?**
We are applying AutoQC, which was also developed in the MacCoss laboratory, as a tool to facilitate longitudinal and rapid tracking of system suitability as one part of a cohesive framework to evaluate the process, system, and quantitative analysis. AutoQC and its predecessor, Statistical Process Control in Proteomics (SProCoP), were developed and implemented by Dr. Michael Bereman and the teams that work on Skyline and Panorama to incorporate statistical process control and automated longitudinal system tracking in proteomics. For more details on AutoQC, please refer to the "Quality control with AutoQC" page on Panorama and the original publication:

Bereman MS, Beri J, Sharma V, Nathe C, Eckels J, MacLean B, MacCoss MJ. An Automated Pipeline to Monitor System Performance in Liquid Chromatography-Tandem Mass Spectrometry Proteomic Experiments. J Proteome Res. 2016 Dec 2;15(12):4763-4769. doi: 10.1021/acs.jproteome.6b00744. Epub 2016 Oct 4. PMID: 27700092; PMCID: PMC5406750.

Beyond peak areas, retention times, and mass error, we suggest examining ratios of precursor area to transition area to delineate between parts of the MS system which may fail. If you will be viewing and processing your data in Skyline, we also frequently examine the idotp (isotope dot product) value to

3. **Why do you recommend dedicated system suitability samples and associated runs?**
   In short, because if a system is not functioning well, then none of the other controls included in your samples are going to be decipherable. You cannot evaluate sample preparation if the measures used to assess it are highly variable due to poor system functioning. The same is true for evaluating batch effects and the impacts of normalization on quantitative results.

   Explicitly evaluating your system can lend greater confidence to your interpretation of other controls. If the system is running consistently with minimal variation, we can be more confident in interpreting variation in our results as being derived from sample preparation or biological variability. We advocate for monitoring of a system as both a diagnostic tool and to minimize more extended system declines.

4. **Why are system suitability runs necessary in addition to the internal QCs when they can often be used to identify problems with the LC-MS?**
   Mostly due to the reasons in Question 3 above. We advocate that if a system is not functioning well, an operator cannot reasonably infer information from any other controls they have included since interpretation will be dependent upon the functioning of your system.

   Without a known sample being measured repeatedly, it would be difficult to determine if a system is functioning solely based on internal QCs in potentially biologically disparate samples. Especially in the context of longitudinal analysis (Say between experiments, which may be months apart by necessity).

   Internal QCs can facilitate earlier identification of some sample preparation and system issues, which is why we insist on including them in all samples. However, we wanted an approach to evaluate our LC-MS systems that could be quickly, easily, and consistently applied across multiple often disparate instruments and be completely independent of the type of sample we were analyzing in a given experiment. A dedicated system suitability QC sample and instrument run will avoid complicating interpretations about the system function due to matrix effects which may change from project-to-project.

   Our logic for this comes in large part from the diverse sample types our group processes each year. In 2023 alone, our projects were run on 6 different MS systems, 3 kinds of LC systems, and in >30 different sample types spanning tissues, cells, biofluids, and synthetic proteins.

   **For example:** How would one assess system performance with internal QCs in disparate sample matrices like plasma, cells, and adipose tissue, which may be complicated by ion suppression or retention time shifts of the quality control peptides? That would affect how we interpret the status of the system. When you process multiple sample types, dedicated system suitability samples and runs provide an unbiased measure of system performance over time that is not impacted by differing sample matrices.

5. **What additional information do external QC samples provide that we cannot glean from internal QCs and system suitability?**
   These external QC samples serve as “known unknowns”. The same pooled external QC sample is processed repeatedly in the same workflow alongside experimental samples. The variation within these technical replicates is indicative of the sample preparation protocol which is also going to affect experimental samples. We use this to evaluate sample preparation reproducibility and the impact of normalization and batch correction on variance. Typically, we do so by evaluating the distribution of the coefficient of variance (CV) of all peptides and, where applicable, all proteins after aggregating peptide signals for a protein-level value.

   We would not expect a high CV from the same sample being processed repeatedly, but processing multiple batches can and typically does increase variation. Additionally, external QCs can be used to evaluate whether the precision of our broader dataset has been improved or not after signal processing. For example - is the normalization and batch correction workflow reducing the variance of these controls? We would expect so if the data analysis is well-selected!

6. **Can the described controls be used to rescue and re-incorporate a sample that suffered from poor or incomplete digestion?**
   In instances where a single sample digestion had failed (For example, if trypsin was not spiked into a single well of a 96-well plate by mistake), typically we would argue that it is not worth the sample preparation time and instrument time to reprocess a single sample.

   However, in scenarios where multiple critical samples failed, or where one batch as part of a larger study suffered a digestion failure, our preferred choice would be to reprocess the entire sample batch. If the same external QC samples we have described in the manuscript was processed in every batch and then processed again for the reprocessed batch, you could determine whether the reprocessed batch is similar to the previously processed samples.

   We would advocate for a minimum of triplicate of each of these external QC samples to be processed when experimental samples are reprocessed. Typically, we have one of each external QC sample processed in a row of a 96-well plate (10 samples, 2 controls).

7. **What other reasons do you have to recommend targeted approaches such as PRM to track system suitability as opposed to an identification-based metric as described in Figure 3’s example with DDA?**
   Anytime database searching and multiple operators are involved, there is always the potential for software updates and setting changes. When multiplied over months or years, this can affect how the system suitability data is interpreted independently of the system function. Targeted acquisition removes the concern about missing data possible in DDA and better facilitates evaluation of the chromatography stability, system mass error drift, peak shape, and overall reproducibility of the method.

8. **Can you summarize what you take into consideration when batching and blocking your samples?**

Other groups have reviewed best practices for sample batching and block randomization in proteomics, and we strongly recommend reviewing their work to determine the best approaches for your experiments.

Burger, B.; Vaudel, M.; Barsnes, H. Importance of Block Randomization When Designing Proteomics Experiments. J. Proteome Res. 2021, 20 (1), 122–128. https://doi.org/10.1021/acs.jproteome.0c00536. PMID: 32969222; PMCID: PMC7786377.

Čuklina, J.; Lee, C. H.; Williams, E. G.; Sajic, T.; Collins, B. C.; Rodríguez Martínez, M.; Sharma, V. S.; Wendt, F.; Goetze, S.; Keele, G. R.; Wollscheid, B.; Aebersold, R.; Pedrioli, P. G. A. Diagnostics and Correction of Batch Effects in Large-scale Proteomic Studies: A Tutorial. Mol. Syst. Biol. 2021, 17 (8), 1–16. https://doi.org/10.15252/msb.202110240. PMID: 34432947; PMCID: PMC8447595.

Briefly, we take all experimental samples and external QC samples and batch them randomly and ensure that they are balanced based on the experimental covariates with manual changes. As an example in biological studies, we ensure that samples are spaced within a batch (or on a plate) such that individuals of different sex, age, and treatment group are not processed in close proximity to one another nor heavily skewed within a batch. For example, we try to ensure that if our experimental cohort is roughly 1:1 for male and female individuals, our plates include individuals representative of as close to a 1:1 ratio as possible. We do the same for other key covariates. We maintain the order in which samples were processed within plates or batches as the run order on the instrument rather than randomizing the samples a second time.
